# Connectomic reconstruction of a cortical column

**DOI:** 10.1101/2024.03.22.586254

**Authors:** Meike Sievers, Alessandro Motta, Martin Schmidt, Yagmur Yener, Sahil Loomba, Kun Song, Johannes Bruett, Moritz Helmstaedter

**Affiliations:** Department of Connectomics, Max Planck Institute for Brain Research, Frankfurt, Germany

## Abstract

The cerebral cortex of mammals has long been proposed to comprise unit-modules, so-called cortical columns. The detailed synaptic-level circuitry of such a neuronal network of about 10^4^ neurons is still unknown. Here, using 3-dimensional electron microscopy, AI-based image processing and automated proofreading, we report the connectomic reconstruction of a defined cortical column in mouse barrel cortex. The cortical column appears as a structural feature in the connectome, without need for geometrical or morphological landmarks. We then used the connectome for definition of neuronal cell types in the column, to determine intracolumnar circuit modules, analyze the logic of inhibitory circuits, investigate the circuits for combination of bottom-up and top-down signals in the column and the specificity of bottom-up and top-down cortical input, search for higher-order circuit structure within homogeneous neuronal populations, and estimate the degree and symmetry of Hebbian learning in the various connection types. With this, we provide a first column-level connectomic description of the cerebral cortex, the likely substrate for a synaptic-level mechanistic understanding of sensory-conceptual integration and learning.

## INTRODUCTION

In neuronal networks as complex as mammalian brains, the existence of repeated circuit modules is the only justified hope for understanding this networks’ operations (Koch 2012). Indeed, the parcellation of the mammalian cerebral cortex into vertically oriented units of about 10^4^ neurons, called cortical columns, was demonstrated for macaque primary somatosensory cortex (Mountcastle 1957), cat visual cortex (Hubel and Wiesel 1962), rodent somatosensory cortex (Woolsey and Van der Loos 1970), and postulated for higher cortical areas, as well (Felleman and Van Essen 1991, Harris, Mihalas et al. 2019). Based on decades of anatomical and functional experiments, a number of main connectivity streams into and within cortical columns have been identified (Gilbert and Wiesel 1979, Gilbert and Wiesel 1983, Jones 1986, Johnson and Burkhalter 1996, Feldmeyer, Egger et al. 1999, Svoboda, Helmchen et al. 1999, Feldmeyer and Sakmann 2000, Lubke, Egger et al. 2000, Schubert, Staiger et al. 2001, Brecht and Sakmann 2002, Brecht and Sakmann 2002, Callaway 2002, Feldmeyer, Lubke et al. 2002, Staiger, Masanneck et al. 2002, Brecht, Roth et al. 2003, Lubke, Roth et al. 2003, Schubert, Kotter et al. 2003, Silver, Lubke et al. 2003, Brecht, Krauss et al. 2004, Brecht, Schneider et al. 2004, Bureau, Shepherd et al. 2004, Feldmeyer, Roth et al. 2005, Shepherd, Stepanyants et al. 2005, Shepherd and Svoboda 2005, Bureau, von Saint Paul et al. 2006, Feldmeyer, Lubke et al. 2006, Schubert, Kotter et al. 2006, Helmstaedter, de Kock et al. 2007, Lübke and Feldmeyer 2007, Sarid, Bruno et al. 2007, Frick, Feldmeyer et al. 2008, Helmstaedter, Staiger et al. 2008, Luo, Callaway et al. 2008, Helmstaedter, Sakmann et al. 2009, Holtmaat and Svoboda 2009, Petreanu, Mao et al. 2009, Sato and Svoboda 2010, Mao, Kusefoglu et al. 2011, Marx and Feldmeyer 2013, Rah, Bas et al. 2013, Koelbl, Helmstaedter et al. 2015, Sarid, Feldmeyer et al. 2015, Yu, Gutnisky et al. 2016, Leinweber, Ward et al. 2017, Feldmeyer, Qi et al. 2018, Winnubst, Bas et al. 2019, Yu, Hu et al. 2019, Wu, Sevier et al. 2023, Huang, Wu et al. 2024). In particular, the sensory input via thalamocortical afferents (Meyer, Wimmer et al. 2010, Meyer, Wimmer et al. 2010, Meyer, Wimmer et al. 2010, Wimmer, Bruno et al. 2010, Oberlaender, de Kock et al. 2012, Oberlaender, Ramirez et al. 2012) into cortical granular layer 4, transmitted to supragranular layers 2/3, from there to infragranular layer 5, and then to distal targets, was described as the canonical circuit (Douglas, Martin et al. 1989) with substantial supporting evidence (Douglas and Martin 1991, Lubke, Roth et al. 2003, Binzegger, Douglas et al. 2004). However, the detailed synaptic architecture of the cortical columnar module is not known. Do cortical layers impose as connectomic clusters? Are layers within the column further structured at the circuit level? What types of networks are implemented among otherwise homogeneous neuronal populations? What is the level of circuit precision within and across cortical layers? Are the so-far considered layers sufficient to capture the main connectomic modules within a column? How is the integration of bottom-up and top-down information implemented at the circuit level?

But even at the level of the cellular components of cortex, key questions remain open. While a certain consensus has been assumed for the definition of types of excitatory neurons, a similarly clear classification of inhibitory interneurons in the cerebral cortex is still elusive (Gonchar and Burkhalter 1997, Kawaguchi and Kubota 1997, Ascoli, Alonso-Nanclares et al. 2008, Tremblay, Lee et al. 2016, Mayer, Hafemeister et al. 2018, Yuste, Hawrylycz et al. 2020, Mao and Staiger 2024). Even an at-scale detailed unbiased sampling of interneuronal type prevalences has not been achieved. Would, however, a connectomic mapping of the circuit context of interneurons allow a clearer definition? Could connectomic information alone allow the classification of neurons in cortex?

And could approaches to extract the degree of learning in cortical networks (Motta, Berning et al. 2019, Dorkenwald, Turner et al. 2022), based on earlier work (Bartol, Bromer et al. 2015), be used to determine hot spots of plasticity within the cortical network?

By acquiring the connectome of a defined cortical column in mouse barrel cortex, we aimed to address these questions. Previous efforts at dense cortical connectomic analyses had provided connectomes at the level of 10^2^ neurons ((Motta, Berning et al. 2019, Dorkenwald, Turner et al. 2022) (Schneider-Mizell, Bodor et al. 2024)). To address the connectomic architecture of a cortical column as a defined circuit module, however, data at the scale of 10^4^ neurons is required, with ideally a circuit-level definition of the bounds of the columnar module. This is relevant both for the definition of modularity, but also for the determination of prevalence for especially rare subtypes of neurons (which, if as rare as 0.01% (i.e. 10^-4^), require the scale of reconstruction to be at 10^4^). We provide such a columnar connectomic reconstruction in the following.

## RESULTS

We obtained a bioptic sample from the primary somatosensory cortex of a 28-day old male mouse centered on the expected location of vibrissal representation (“barrel cortex”, (Woolsey and Van der Loos 1970) Fig. 1A). Location within barrel cortex was confirmed by cytochrome oxidase staining of the remaining tissue (our sample contained parts of barrel rows C and D). The sample was stained and embedded using the Hua protocol (Hua, Laserstein et al. 2015), followed by ultrathin cutting in combination with automated section pickup using ATUM (Hayworth, Morgan et al. 2014). Imaging was performed using a 61-beam multiSEM (Zeiss, Oberkochen, Germany, (Eberle, Mikula et al. 2015), Fig. 1B). With this, we obtained a dataset sized 1.3 x 1.3 x 0.25 mm^3^ at 4x4x35 nm^3^ voxel size (7,199 consecutive sections, cut over ∼28 hours, and imaged using 50 ns dwell time per pixel over total of 64 days, effective throughput 18 Tvx/day, methods similar to (Loomba, Straehle et al. 2022), Fig. 1C). After alignment, the dataset was processed using automated segmentation and agglomeration methods (conceptually similar to (Motta, Berning et al. 2019), but in an improved form implemented as voxelytics package by scalable minds, Potsdam, Germany). Automated synapse, blood vessel, cell body and myelin detection were performed (Fig. 1D). Based on overall cell densities, the likely location of barrels in L4 was determined, and a bounding box sized 360 µm in width and 1.3 mm in height was defined for further processing, centered on a presumed barrel. Then, thalamocortical afferents were identified (using previously established criteria about synapse density and multiple-target boutons, (Bopp, Holler-Rickauer et al. 2017, Motta, Berning et al. 2019), yielding a map of VPM synapses (Fig. 1E). This map allowed the delineation of the barrels in L4, outlining the presumed barrel columns in mouse S1 cortex. Then, all cell bodies within the column-centered volume were classified into glia, neuron and vessel-related (Fig. 1D). For all neurons, dendritic reconstructions were completed by iterated ending detection, visual inspection, and identification of the split-off dendrite pieces (total ∼10,000 work hours for total of 30 meters of dendrites; note these work hours were required on an earlier segmentation version and are dispensable in the latest version). Then, spine heads were automatically detected (84.6 million) and attached to the corresponding dendritic shaft using automated neurite flight reconstruction (RoboEM, (Schmidt, Motta et al. 2024)), yielding dendritic reconstructions with 97% recall and 98% precision (for dendritic spines: 94% recall and 97% precision). For reconstruction of axons, automated ending detection followed by RoboEM-based automated flight continuation was used to extend axons by several millimeters per neuron (see (Schmidt, Motta et al. 2024)). For inhibitory interneurons (INs), major remaining axonal mergers were resolved after visual inspection (about 100 work hours). With this, we obtained the reconstruction of n=92 million synapses and n=9500 neurons in the column-centered volume (Fig. 1J), yielding a 9500x9500 connectomic map with 11.47 million mutual synapses (Fig. 1I).

**Figure 1:**
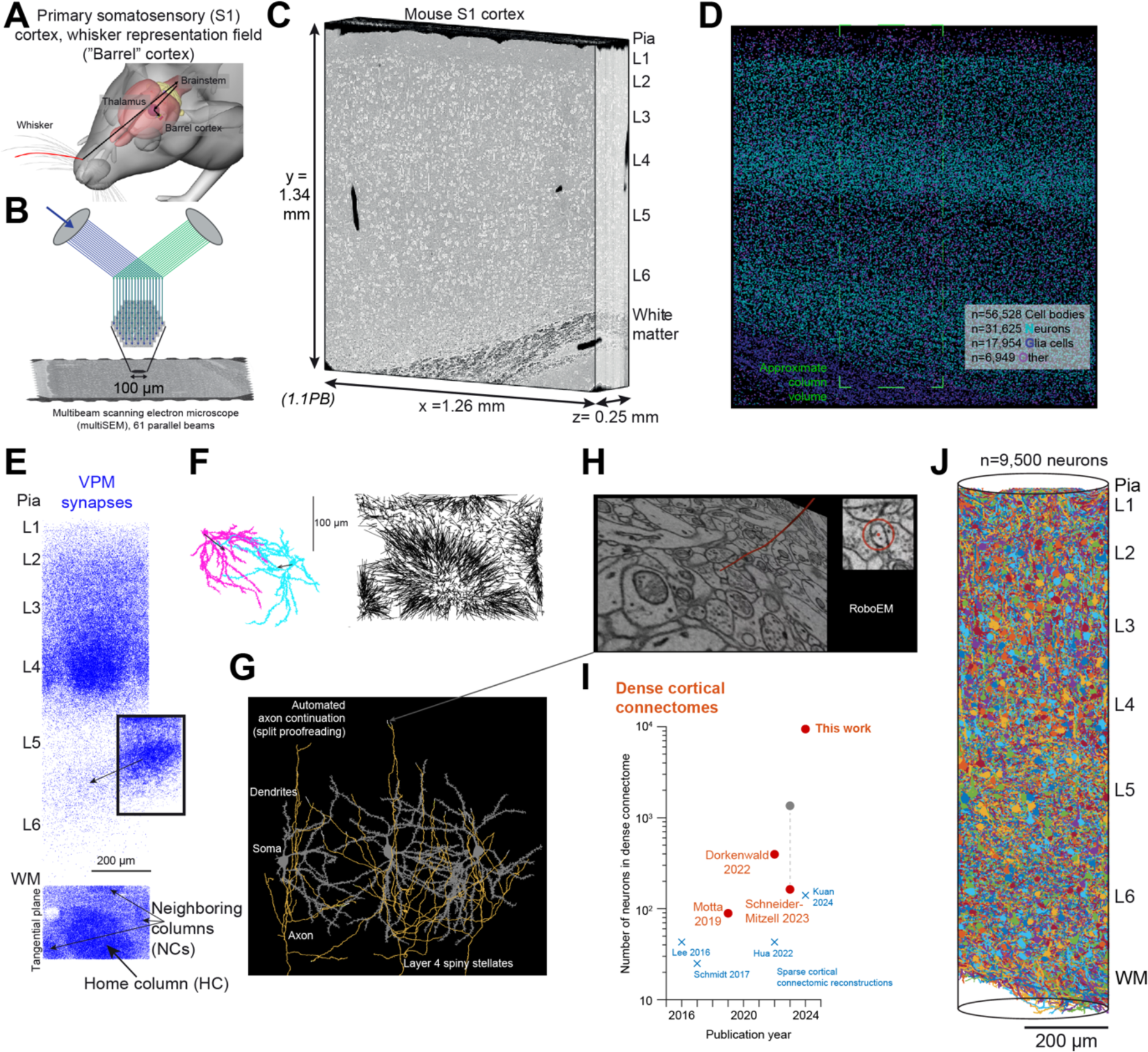
Connectomic reconstruction of a cortical column from 3D EM data. **(A)** Whisker-to-barrel system in rodents as one prime example of topographic organization in the nervous system, and mouse primary somatosensory (S1) barrel columns as a prototype of mammalian cortical columnar architecture. **(B)** Acquisition of large-scale 3D EM dataset using ATUM-multiSEM technology (Eberle, Mikula et al. 2015). **(C)** Petascale (1.1 PB) 3D-EM dataset of mouse barrel cortex acquired at 4x4x35 nm voxel size. **(D)** Reconstruction of n=56,528 cell bodies in the dataset, with subvolume centered on one barrel column highlighted (dashed lines). “Other” are primarily blood-vessel related cell bodies. **(E)** Map of VPM synapses (subsampled 20-fold) in the column volume. Coronal view, top shows home column barrel as high-density cluster. Tangential view, bottom, shows home column and three neighboring columns as high-density regions. Inset, high-density VPM at L5B-L6 border shown without subsampling. **(F)** Analysis of dendritic morphologies of excitatory neurons (ExNs) in layer 4 (L4), which have been reported to exhibit strong dendritic asymmetry towards the home column (Feldmeyer, Egger et al. 1999, Lubke, Egger et al. 2000). Two example neurons, left, with mean dendritic directionality vector in the tangential plane, and quiver plot of all dendritic directionality vectors for n=2,002 neurons in L4 showing similar columnar pattern as from VPM synapse distribution (compare F, right with E, bottom). **(G,H)** Examples of L4 spiny stellate reconstructions. Axons were, after initial automated agglomeration, automatically continued using RoboEM automated flight tracing ((Schmidt, Motta et al. 2024), inset in H shows example RoboEM tracing, adapted from Suppl. video of RoboEM paper). Example reconstruction shown was based on an earlier segmentation version. **(I)** Comparison of sparse (crosses) and dense (circles) connectomic reconstructions in cortex. Size of a connectome is minimum of pre- and postsynaptic neurons (i.e. requiring a square connectome involving all connections between all neurons, red). Gray dot: number of postsynaptic neurons, however not as dense connectome. **(J)** Reconstruction of n=9423 of n=9900 neurons in the column volume and their n=11.47 million synapses (out of the 92 million synapses in the column volume), this study.

### Connectomic definition of cortical columns and excitatory neuron types

As a first attempt we investigated the degree to which the connectome alone, without any further information on neuronal morphology or position in the cortex, would allow the definition of a cortical column, and identify relevant excitatory and inhibitory cell types. We first used dendritic spine rate (Fig. 2a) to separate excitatory neurons (ExNs) from inhibitory interneurons (INs). The distributions were clearly separate, with the exception of deeper layer 5. Here, we used the targeting fraction of spines by neuronal axons to distinguish ExNs from INs (see below for the reason for the overlap in L5). Then, we applied a standard clustering algorithm to the full ExN connectome (Fig. 2B,C, Ward’s method applied to the ExN-ExN input connectome as data matrix). In fact, the resulting connectomic clusters yielded very clear delineations of cortical layers and barrels in L4 (Fig. 2D,E). Not only the centered home column, but also the three neighboring columns, with only fractional coverage in the dataset, were delineated clearly based on the connectomic clustering (Fig. 2D). More specific geometric arrangements of neurons were recovered by stepwise cluster decomposition (see Methods): the spherical shape of the L4 barrel (Fig. 2E,G), the corresponding delineation of L5A as a curved arrangement (Fig. 2E,G), the subdivision of L4 into a lower and upper part with populations of spiny stellates (sps) and star pyramidals (stp) in each (Fig. 2E-G; extending the depth-based division found in (Hua, Loomba et al. 2022)). Furthermore, the clustering indicated a separation of the commonly lumped-together “L2/3” into L2, L3A, L3B, L3C. In addition, unexpectedly, there was a separation into a column-centered L2 cluster, and the surrounding L2 cells, which often showed a slanted apical dendrite orientation towards the column center (Fig. 2F). L5A could be separated into a pool of L5A cells that aligned with the bottom barrel border in similar curvature (Fig. 2E,G), followed by more classical L5A cells. Then, a subtype of L5 cells was separated that had large but sparse dendritic trees, and a low spine rate (the reason for the apparent overlap of spine rates in L5, see Fig. 2A; see also (Schneider-Mizell, Bodor et al. 2024) for this subtype). L5B cells were identified, and a very clear separation of L6 ExNs into those in L6A with an apical dendrite reaching into L4, and L6B with apical reaching to L5A, but not L4 (Fig. 2G). With this, 17 ExN types were directly a result of the connectomic clustering.

**Figure 2:**
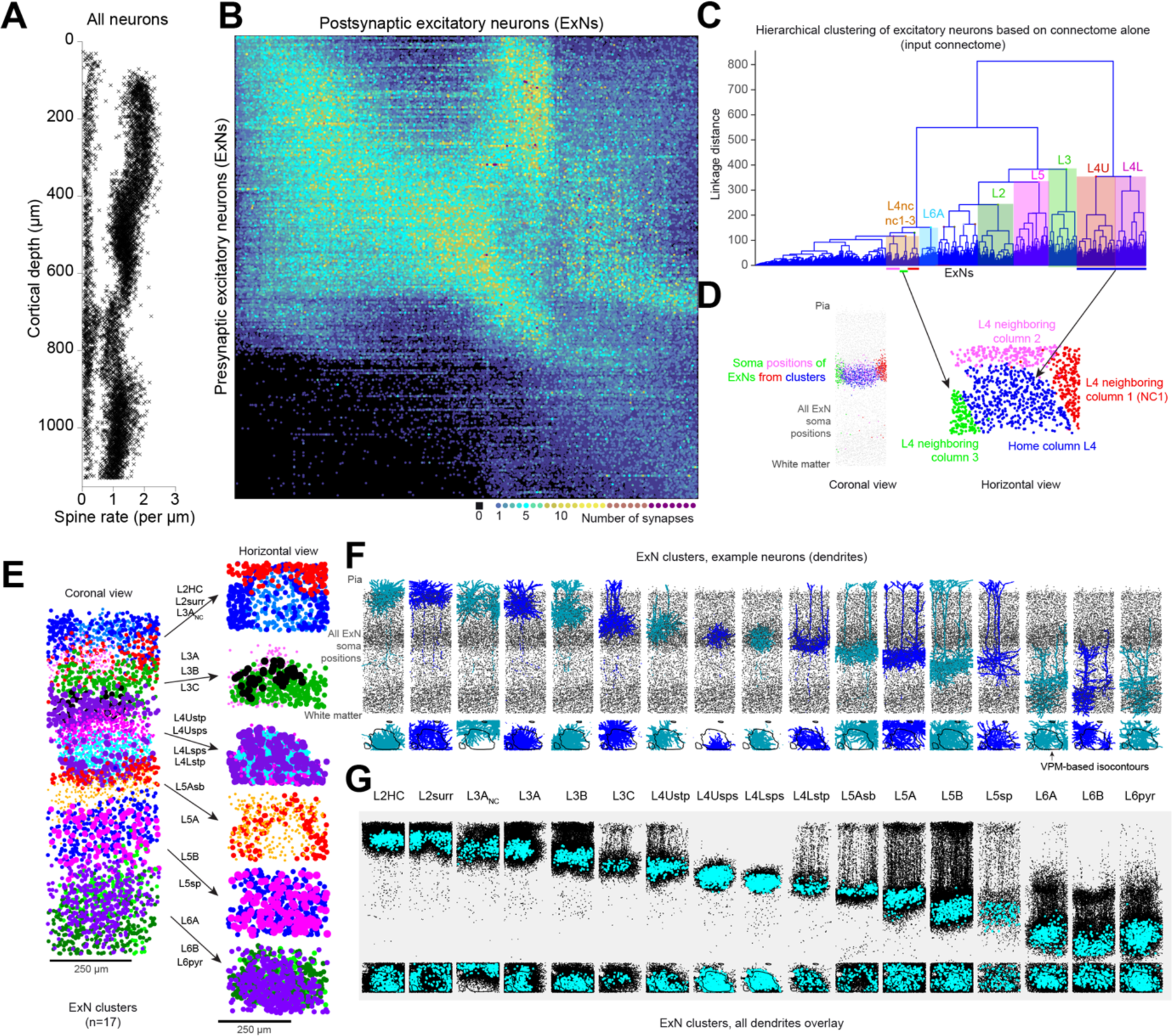
Connectomic definition of a cortical column and its layers, and of excitatory neuronal cell types (ExNs). **(A)** Depth distribution of rate of spines on dendrites of individual neurons. Separation into excitatory (ExNs, spiny) and inhibitory (INs, less spiny) neurons clearly visible for all depths except a L5-related region. In this region, rate of axonal output onto spines was used as second criterion to separate ExNs vs INs. **(B)** Connectome of all ExNs, used for clustering and ensuing identification of ExN types. Note that while this matrix is sorted by soma depth, the ensuing clustering was performed only on the input connectome (B), thus no explicit depth information, cell morphology or other priors were used, only the input connectivity per ExN from all other ExNs. Input connectivity was used since it was assumed to be less affected by variable axon completeness across ExNs. Input from INs was not used so E-I and I-E connectivity could be later used as a corroboration of identified cell types. Earlier version of ExNs shown with n=6655 neurons, local outdegree outliers removed. Note main cortical connectivity is already visible, with upwards and downwards signal flow, including sparser L5->L2 and L3 connections; note however that this picture needs to be refined when cell type prevalence and synaptic multiplicity are considered, see Fig 6. **(C,D,E)** Hierarchical cluster analysis (Ward’s method) of ExN-ExN connectome shown in B. The barrels in L4 for the home column and 3 neighboring columns are readily identified as first main cluster separations, as are the main cortical layers. Note that no depth or morphological information was used for this cluster analysis, showing the power of connectomic definition of cortical columns and main cortical layers (see E). **(E)** Main ExN cluster separations across cortical depth (right, top to bottom): center-surround structure in L2, separation of L2 and L3, sublayers of L4 (see F,G), center-surround structure in L5A below the barrel; columnar clustering in L6 indicating extent of columnar order across all cortical layers (L2 – L4 – L6). **(F,G)** Examples of ExNs per ExN type (total of 17 ExN types, of these 15 HC-related) and overlay of all ExNs (G). Cyan dots: cell body locations for each type. Note again that depth relations are result of cluster analysis, not explicit layer boundary definitions. Clusters were however refined for depth homogeneity, see methods, affecting a small subset of neurons. Abbreviations in ExN type names: HC, home column; surr, surround; NC, neighboring column(s); L4Ustp: L4 upper star pyramidals; L4Usps: L4 upper spiny stellates; L4Lsps: L4 lower spiny stellates; L4Lstp: L4 lower star pyramidals. Note that the subdivision of L4 in upper and lower was a result of the clustering (see Methods), not a preconceived separation (cf. (Hua, Loomba et al. 2022) where stp and sps, but not sps within were found to be separated). Gray dots in F: all ExN somata in the dataset. Contour lines in F, bottom panels: isocontours of VPM synapse densities (from data in Fig. 1E). Note that color codes are not consistent across panels.

We next used a similar approach for INs, and also asked whether the here-found subdivision of ExN populations would be matched by IN target specificities, which would support this connectome-only based cell type definition further.

### Connectomic definition of interneuron subtypes

We first clustered the IN connectome (input from all neurons and output to all neurons as clustering dimensions, n=18000 dimensions) to obtain a first rough IN assignment. Indeed, this already revealed the barrel-related inhibitors (Koelbl, Helmstaedter et al. 2015, Feldmeyer, Qi et al. 2018) in L4, sublayer-specific inhibitors of L5A, L5B, L6A and L6B, as well as layers 3 and 2. Furthermore, many L1 INs were assigned to common clusters. Importantly, INs commonly described as “bipolar” or “bitufted” were largely separated into different clusters at this initial step. We then incorporated 5 additional connectomic parameters, that have been previously shown to distinguish IN types (Fig. 3C): the ratio of output synapses made onto other INs (IN output fraction), the fraction of inhibitory input received by the IN (i/(i+e) ratio), the fraction of synapses made onto postsynaptic cell bodies (at multiplicity 2), the rate of input spine synapses along the IN dendrites, and the fraction of input synapses made by VPM. These were used as additional separating parameters.

**Figure 3:**
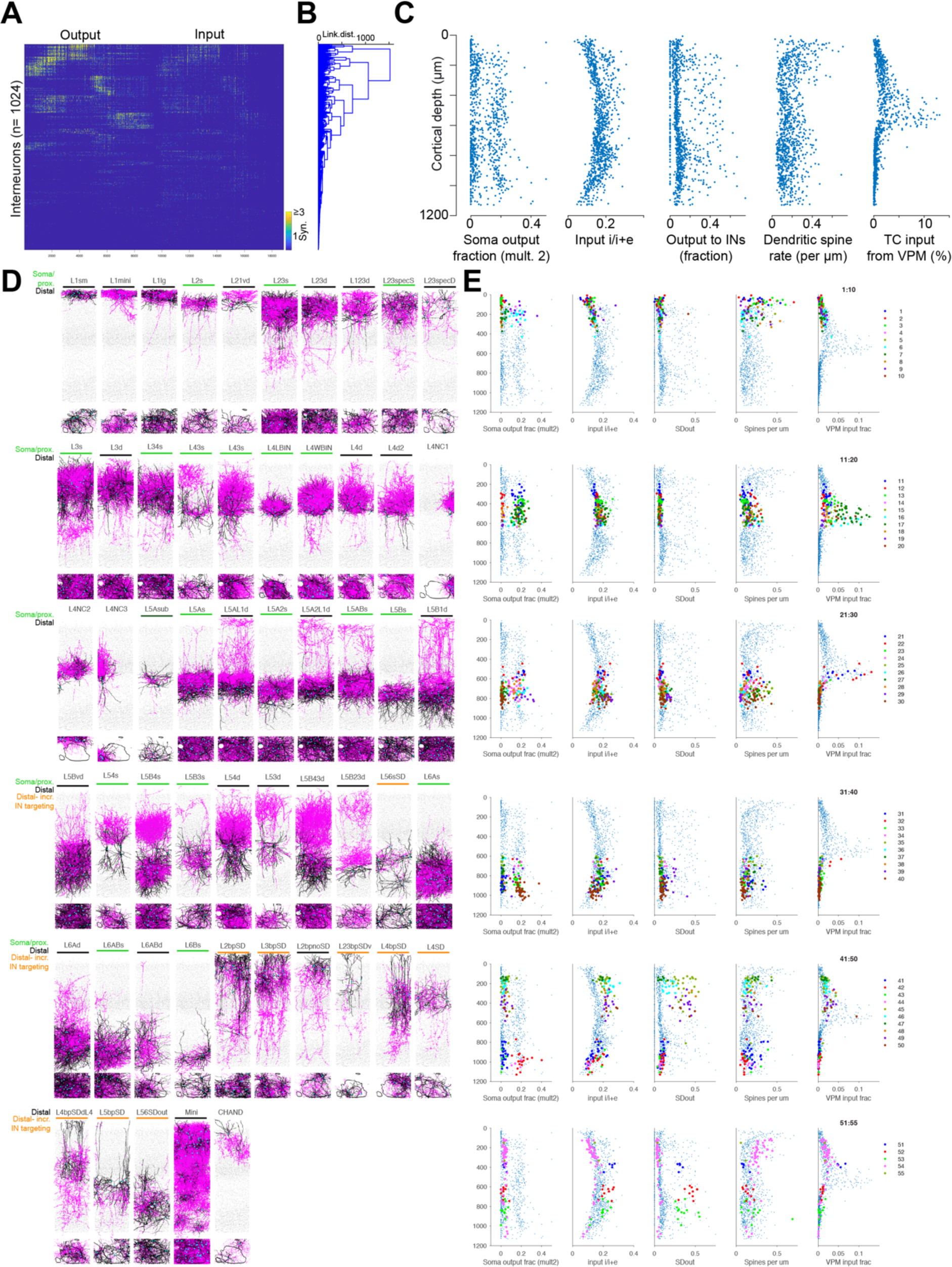
Connectomic definition of IN types. **(A)** Connectomic data matrix used for IN type definition: n=1024 INs (sorted by soma depth, but irrelevant for the ensuing cluster analysis) times their output to all neurons, concatenated with their input from all neurons (data matrix of size 1024 x 19000). Here, in contrast to clustering of ExN connectome, connectomic interaction between IN and ExNs was used for clustering, since it had to be assumed to be essential for IN subtype definitions. **(B)** Hierarchical clustering (Ward’s method) followed by separation of clusters according to 5 additional connectomic parameters **(C)**: Output to somata (only connections with at least 2 synapses considered); input i/(i+e) balance; Fraction of output to INs (out of all output synapses); spine rate along dendrite, fraction of input from VPM. Note that while only connectomic parameters (A and C) were used for quantitative cell type definition, the plausibility of cell type separations was qualitatively judged, which included IN morphology as a qualitative criterion. **(D,E)** Resulting n=55 IN types, of which some are very low prevalence (e.g. n=2 Chandelier neurons). Dendrites, black; axons, purple. Soma/proximal vs. distal innervation preference as well as preference for IN innervation shown as colored lines. Parameter plots in E show soma output as main additional connectomic separation criterion, with input i/(i+e) relevant for bipolar cell types (last but one row), and output to other interneurons (last two rows). Note also homogeneous clusters in spine rate and VPM input for some IN types.

First we inspected the L4 INs, which had previously been shown to comprise barrel-related INs(Koelbl, Helmstaedter et al. 2015), which were suspected to be specific to spiny stellate vs star pyramidal ExN targets based on a small connectomic sample (Hua, Loomba et al. 2022). Indeed, clear clusters emerged in which barrel-INs were clustered together, which were further differentiable by their lower-barrel vs. whole- barrel targeting (L-BIN vs W-BIN, (Hua, Loomba et al. 2022), Fig. 2D). Importantly, even though we had only partial reconstructions of the neighboring barrels, separate IN clusters comprised those INs in the neighboring barrels that had the same BIN, L- Bin and W-BIN characteristics (Fig. 2D). Notably, these INs were soma-preferring, as previously documented (Fig. 3E), did not substantially innervate other INs, and received the highest proportion of VPM input synapses of all INs (Fig. 3E). In addition, we found two barrel-related IN types with preference for distal dendrite innervation (Fig. 3D,E).

Next we noted a small-dendrite large-axon IN in several supragranular clusters, which turned out to have a high rate of spiny input synapses. By separating high-spine rate INs, we identified this IN type to be present mostly in L1-L4 (Fig. 3D,E, “Mini”IN type). This IN type appeared to be arranged in a dendritic mosaic, tiling the L4-L2 volume. The IN was very sparse below L4. It had morphological similarities to a recently described IN class (Machold, Dellal et al. 2023).

Further, the clustering identified the large soma-preferring INs with axons specific to L2, L3, the surrounding of barrels in L4, L5 and L6 (Fig. 3D). L1 INs were of mainly three types: those of the mosaicking IN described above (“mini”), a larger IN type, and a smaller type (Fig. 3D, L1sm, L1mini, L1lg).

In terms of INs with multi-layer axons, the most notable were the L5B-L4 INs (Fig. 3D, type L5B4s) and the L5-L1 doubly innervating INs (likely Martinotti cells, types L5AL1d, L5A2L1d, L5B1d in Fig. 3D, (Wu, Sevier et al. 2023)).

Vertically oriented (“bipolar”) INs were identified by higher i/i+e input (located mainly in L2 and 3, Fig. 3E, see (Loomba, Straehle et al. 2022). Some of the most often-studied IN subtypes were the rarest: the dataset contained n=2 Chandelier neurons in L2 (Fig. 3D, CHAND type). This is important, since it sets a lower bound on the Chandelier IN prevalence in relation to relevant cortical modules, for which both the definition of a cortical module and the required scale of reconstruction were necessary (10^4^ with 10^3^ INs). The low rate of Chandelier INs (about 1-3 per column, i.e. about 10^-4^ or 0.01%) also sets limits to their potential encoding capabilities; and their ability to carry sensory specific vs. global signals. For further implications of IN prevalence, see also Fig. 7C.

### Inhibitory effects on excitatory signaling in the column

We next investigated how the various types of INs are recruited in the excitatory processing in the columnar circuit (Fig. 4). For this, we first considered the homotypic ExN-ExN connectomes within layers (Fig. 4A), which showed connectivity rates of 5- 35% (Fig. 4B), with most connections made via 1 synapse per pair (corroborated by manual analysis of a subset of spiny stellate neurons), and reciprocity rates at the level expected from the pairwise connectivity rates (Fig. 4B).

**Figure 4:**
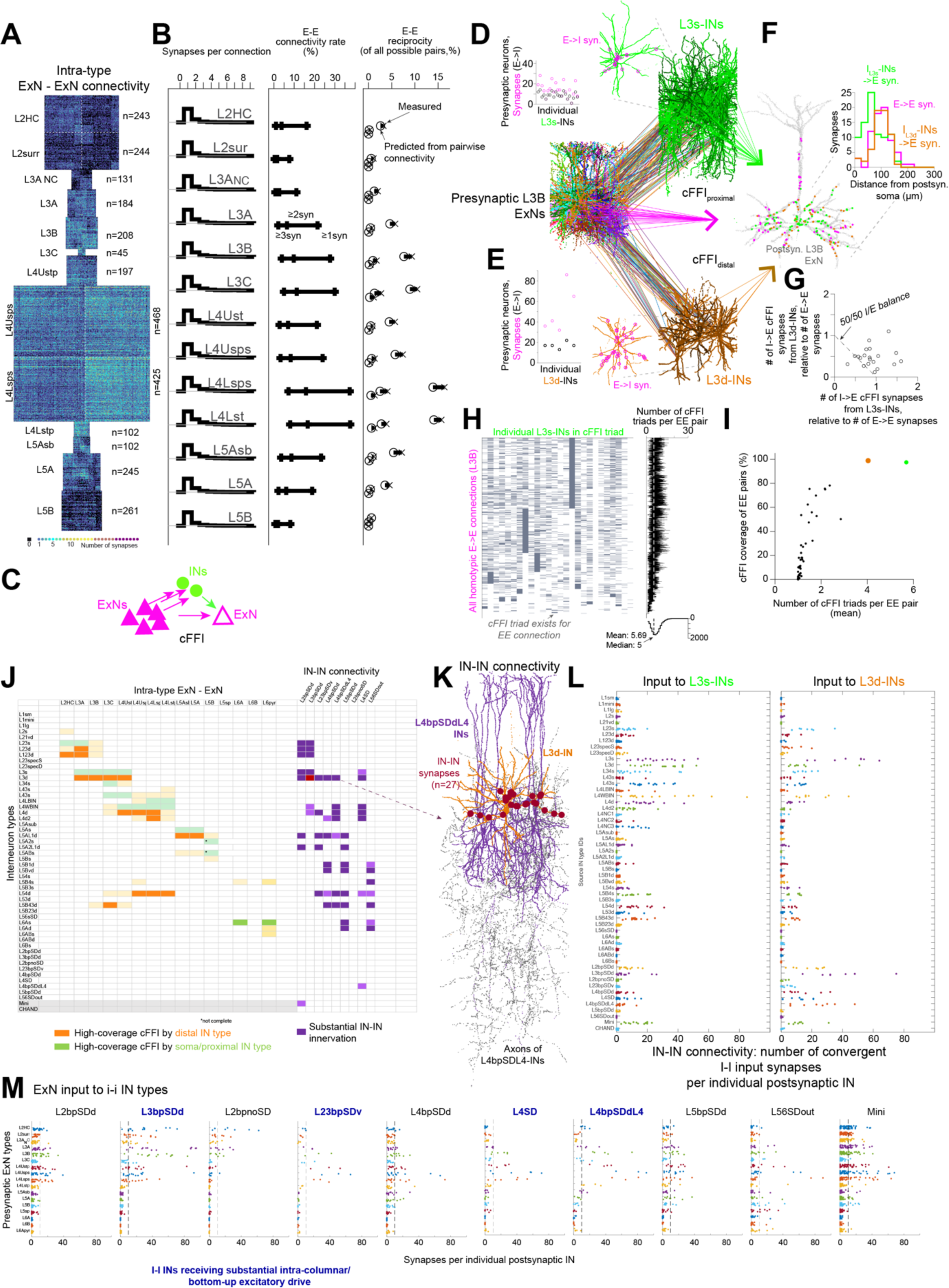
Connectomic analysis of homotypic feedforward processing, and its inhibitory and disinhibitory components. **(A)** Homotypic ExN-ExN connectomes for types of ExNs in the column. (B) Synapses per unitary connection, connectivity rate and reciprocity for connections with at least 1, 2 and 3 synapses shown. Note most frequent number of synapses per connections was 1 (similar to the results from a ground-truth L4 connectome, (Sievers 2023)). Note further that reciprocity is not exceeding the prediction from pairwise connectivity for any connection. **(C)** Sketch of cellular feedforward inhibition circuitry: the pool of ExNs that strongly drives a given postsynaptic ExN is also innervating populations of INs that in turn innervate that very same postsynaptic ExN. In such circuits, depending on their precision of synapse placement, axon diameters and convergence, effective control of signal propagation including shutdown of processing can be achieved (Schmidt, Gour et al. 2017). **(D-F)** Example feedforward connection: all L3B ExN inputs to a given L3B ExN shown (left). The INs that this presynaptic ExN pool innervates strongly are L3-related soma- innervating INS (L3s-INs, D), and a second population of L3-related distally- innervating INs (L3d-INs, E) that were even more strongly innervated by the ExNs. Together, these three presynaptic populations (L3B ExNs, L3s-INs, L3d-INs) innervate the given L3B pyramidal cell (F). When comparing the number and spatial distribution of these synaptic pools in the cellular feedforward inhibition triads, the distal inhibitory synapses match the excitatory positions, and are at least as numerous as the excitatory inputs. **(G)** summary of cFFI inhibitory-excitatory balance on postsynaptic ExN dendrites shows very strong inhibitory bias, reaching and exceeding 50%. Since the E->I activation was also very substantial (see C, D), this circuit provides a very strong cFFI suppression of postsynaptic activity (see (Schmidt, Gour et al. 2017) for detailed biophysical modeling of such circuits). Note this analysis assumes sparse neuronal activity, such that at a given point in time, only a subset of a neuron’s presynaptic ExN populations are synchronously and briefly active. **(H)** Systematic cFFI in L3B-L3B ExN-ExN connectome: all ExN-ExN connections with at least one synapse (vertical axis), and INs of the 3s type (horizontal axis) shown; gray indicates that the cFFI triad for this particular E-E pair and individual IN was found. Right, average number of cFFI triads per EE pair is 5.7, indicating strong cFFI. Note that almost all EEpairs are receiving multiple cFFI. **(I)** Quantification of cFFI strength and coverage for L3B-L3B ExN-ExN connections, shown for all IN types. Note only two IN types show high coverage (>90% of EE pairs) and strong average multiplicity (>4 INs involved in cFFI per E-E connection); these two IN types are L3s and L3d, as shown in C-F. **(J)** Systematic analysis of all homotypic E-E connections in the column (left to right), and indication which INs provide strong and high-coverage cFFI. Note that for each E-E connection type, at least one pair of soma-innervating (green) and distally- innervating (orange) IN type is providing strong cFFI. Lighter shades: weaker contributions. Which INs can shut off these strong cFFI IN types? Right: purple entries indicate which IN-preferring IN types provide substantial inhibition to the various types of cFFI INs. **(K)** example of L4-related bipolar cell type that provides strong inhibition of L3d INs. **(L)** Survey of all inhibitory input to L3s and L3d INs to search for disinhibitory circuits (of which the connection shown in J is one example). Note that connection strength is reported as number of convergent inhibitory synapses per postsynaptic neuron. This is important since prevalence of IN types varies, and strength of unitary connections can compensate for low-prevalence presynaptic pools. Note that bipolar-IN input is strongest for L3d compared to L3s INs, note further that the most substantial input to L3s INs is provided by other s-type and d-type INs that have no overall preference for IN innervation. **(M)** Which ExN types can activate disinhibitory IN types? Analysis of drivers of disinhbitory INs. Note that a subset of bipolar-type INs receives strong activating synaptic input from intracolumnar excitatory populations, notably L4 and L3, thus the direct bottom-up input of columnar activation (blue bold labels), in contrast to the picture that disinhibitory INs are primarily driven by top-down input. Other subtypes do receive little intracolumnar excitatory input, suggesting influence form distal sources, see next figures.

We then considered a prototypical ExN circuit, the L3B connectome with about 30% ExN-ExN pairwise connectivity, and asked which IN types were involved in possible cellular feedforward inhibitory circuits (Fig. 3C). For this, we considered a L3B ExN (Fig. 4D-G), and determined its presynaptic L3B ExN input population (Fig. 4D). Then we asked which IN types were recruited by this particular ExN input population. As expected from feedforward inhibitory circuits (Buzsaki 1984, Pouille and Scanziani 2001, Schmidt, Gour et al. 2017), a type of soma-preferring IN was strongly innervated, the L3s-INs (Fig. 4D). These INs in turn projected substantially to the postsynaptic L3B neuron (Fig. 4F) with numerous proximal synapses, sufficient to counterbalance the excitatory inputs received in this particular cellular feedforward inhibition (cFFI) setup (Fig. 4F). To our surprise, however, a second type of L3 INs was recruited by the same presynaptic ExN pool, with even stronger synaptic convergence (Fig. 4E vs 4D). This second IN type, however, targeted the postsynaptic ExN more distally (type L3d INs), matching the spatial distribution of the ExN->ExN synapses onto the postsynaptic L3B ExN (Fig. 4F). Together, the two cFFI circuits provided strong inhibitory input that outweighed the ExN->ExN synapses (Fig. 4G). We next asked whether such cFFI circuitry was present for other intra-layer ExN connectomes, and how systematically these circuits were implemented (Fig. 4H-J).

First, for investigating the coverage of cFFI motives within a given ExN-ExN connectome, we listed all individual ExN->ExN connections within L3B (Fig. 4H) and determined for each individual L3s-IN whether it closed the cFFI triad for each particular synaptically connected E-E pair. This cFFI overlap matrix (Fig. 4H) allowed us to determine the coverage (fraction of E-E pairs accompanied by at least one cFFI triad) and redundancy (number of INs of a given type that provide cFFI per synaptically connected E-E pair) of cFFI for a given IN type and E-E connectome. When applying this analysis for all IN types (Fig. 4I) for the L3B E-E connectome, we found that indeed L3s-INs and L3d-INs are the two types with more than 90% coverage, and on average more than 4 and 5 cFFI-INs per E-E pair, with all other IN types providing weaker and less consistent or no cFFI coverage (Fig. 4I).

We then investigated all homotypic E-E connectomes (Fig. 4J) and determined based on these overlap matrix statistics, which IN types provided high-coverage and substantial cFFI. We found that indeed for each intralaminar connection type, at least one soma-preferring and one distal-dendrites preferring IN type provided substantial cFFI as defined for the L3B case above. We conclude that efficient and strong cFFI is provided systematically by combinations of proximal and distally innervating INs within the column, and that this cFFI is substantial enough to shut down intracolumnar excitatory signal propagation (see Fig. 4G), allowing for highly selective gating (Vogels and Abbott 2009, Schmidt, Gour et al. 2017).

We next asked which of the IN types is in turn able to shut off this strong inhibitory cFFI influence, such as to allow the gating of information in the column (Fig. 4J-L). For this, we first considered the “bipolar” types of INs, which have been associated with providing disinhibition in cortical circuits (Pfeffer, Xue et al. 2013, Kubota, Karube et al. 2016, Yu, Hu et al. 2019, Lukacs, Francavilla et al. 2022). Indeed, we found substantial IN-IN innervation from these IN types to those IN types that are recruited in the cFFI circuits per layer, with differential preference for INs in various layers (Fig. 4J). An example of such innervation is shown in Fig. 4K for the L4bpSDdL4 type providing substantial I-I innervation of a postsynaptic L3d IN.

To obtain an unbiased sampling of I-I innervation for the example of L3s and L3d IN types, we analyzed the number of synapses made onto individual L3s and L3d INs from all types of IN in the column (Fig. 4L). For L3s-INs, while some innervation from bipolar IN types was found (Fig. 4L, left bottom), the quantitatively strongest inhibition was received from other L3s, from L3d, and from several IN types in L4 and L4/L3. This is already notable for several reasons. First, there is not only substantial self-inhibition within L3 that would yield disinhibitory relief after the initial inhibitory activation, but substantial (quantitatively similar magnitude) disinhibition received from INs in the input layer 4. This implies that activation of L4 INs already provides a pre- disinhibition of L3sINs, possibly gating the excitatory signal flow. Secondly, it is noteworthy that the inhibitory input to L3s-INs is quantiatively similar from other L3s (self-inhibition) and from L3d INs. Since L3d INs are only less than half in number (n=11 vs n=26), this strong connection requires a more than 2-fold larger unitary connection strength (in terms of number of synapses per connected pair) for L3d->L3s than vice versa. While analyses focusing on unitary connection strength would conclude a more than 2-fold stronger inhibition in one direction, the inclusion of neuronal prevalence reveals a balanced L3d->L3s vs L3s->L3s inhibition. This is one example of why considering the total number of converging synapses from a presynaptic population is important compared to the analysis of single unitary paired connections. When analyzing the inhibitory input to L3d INs (Fig. 4L, right), their quantitatively strongest inhibitory input originates from L3s INs (with substantially less self inhibition), and from bipolar IN types. The latter is substantial enough (especially from L3bpSDd) with 20-80 synapses per postsynaptic neuron to predict the ability to shut down the L3d-IN-based cFFI.

We therefore finally asked whether excitatory neuronal populations within the column can drive the disinhibitory circuits, or whether they are primarily innervated by possible top-down inputs (Fig. 4M). We found that for a subset of bipolar IN types, intracolumnar excitatory populations provide strong synaptic input, in particular from layers 2+3 and 4. This goes against the circuit picture in which top-down input drives disinhibitory cortical innervation, and shows that already the cortical input layers (and also VPM directly, see below) can drive the “bipolar” IN-IN IN system.

Together, these data suggest a highly recurrent inhibitory-disinhibitory system paralleling a recurrent ExN excitatory network, in which IN types are activated in parallel to ExNs, and propagate inhibitory-disinhibitory dynamics within and across layers. We do not find connectomic evidence for the simplified picture of sequential recruitment of inhibitory subtypes (see e.g. (Yu, Hu et al. 2019)).

### Layer 1 inputs to the cortical column

With the ultimate goal of analyzing the integration of bottom-up (sensory) with top-down information in the cortical column, we next wanted to understand the logic of input within layer 1, which is thought to carry long-range top-down information into the sensory cortex (Fig. 5, (Johnson and Burkhalter 1996, Mao, Kusefoglu et al. 2011, Wang, Sporns et al. 2012, D’Souza, Meier et al. 2016, Winnubst, Bas et al. 2019)). When first counting the number of input synapses made in L1 for all excitatory cell types, L2 ExNs stand out by receiving at least 2-fold the number of synaptic inputs in L1 than any other ExN cell type, even compared to the only slightly deeper located L3A ExNs (Figs. 5A,B). Since synapses within L1 are comprised of long-range axonal inputs and local inputs to layer 1, we next selected axons that were exclusively within L1, traversing L1 without diving into the deeper layers (or originating from there), and were spatially not biased in the horizontal plane (Fig. 5C,D). These axons were further extended using RoboEM. We then asked whether within such a population of traversing L1 axons, targeting biases for the types of ExNs in the column could be observed. When comparing the number of output synapses made onto L2HC ExNs vs those made onto L5A+L5B ExNs (Fig. 5C), we found a certain bias of a subset of axons towards L2HC innervation vs those that were either unbiased or had a slight preference for deeper layer ExNs. We then subdivided this set of L1 axons according to their L2HC vs other innervation preference (Fig. 5D). Fig. 5E shows an example axon targeting 20 L2HC but only 3 L5 ExNs. When we then analyzed the innervation of all ExN and IN types by these two L1 axon populations (Fig. 5F), we found that in addition to the L2HC vs L5 preferences which had been the basis for their subdivision, the axon populations differentially innervated certain IN types (Fig. 5F, right): two of the L1 INs were preferentially innervated by the non-L2—preferring L1 axons. This differential output to INs could be interpreted as a corroboration of the initial L1 axon subdivision.

**Figure 5:**
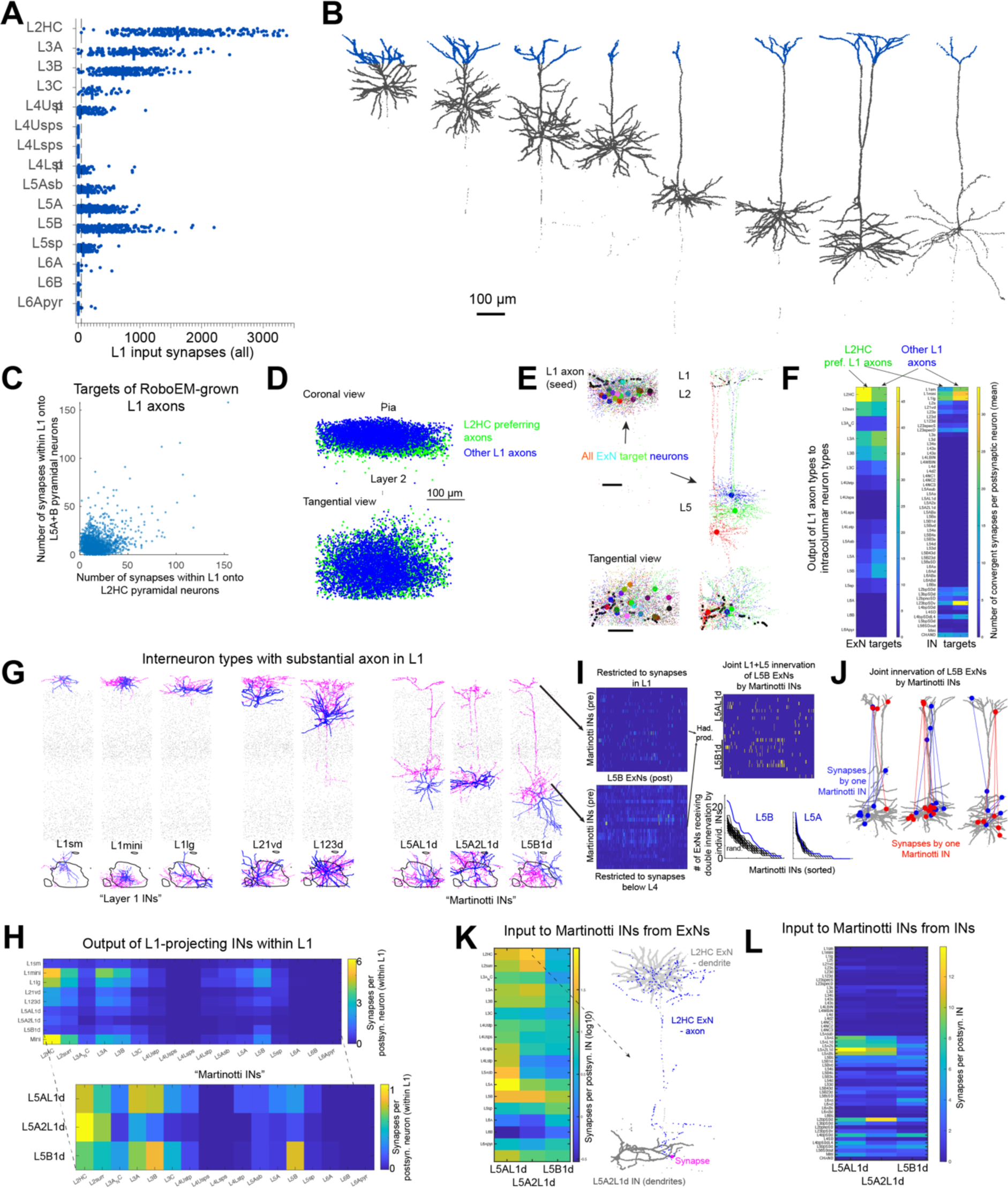
Analysis of L1 input and L1-related inhibitory circuits. (A,B) Number of synaptic inputs within L1 for all ExN cell types (A), examples shown in (B). For simplicity, a depth threshold was used that matched the transition L2->L1. **(C)** Synaptic targeting bias of L1 axons towards either L2 or L5A+B pyramidal neurons, analyzed for a subset of n=662 L1 axons that were not linked to any neuron in the column and were RoboEM-reconstructed to obtain more complete input axons. Note slight bias of a subset of axons to innervate L2 pyramidal neurons. **(D)** spatial location of output synapses of L2-preferring vs. other L1 axons, note slight spatial offset along cortical axis. **(E)** Example of all ExN targets of one L2-preferring L1 axon: 20 L2 pyramidals but only 3 L5 pyramidals targeted. **(F)** Innervation preferences of L2-preferring and other L1 axons. Note that this ExN-targeting based L1 axon discrimination is correlated with IN targeting preferences: non-L2 preferring axons show substantially higher targeting of subtypes of L1 INs, further corroborating the L1 axon type subdivision. **(G)** Analysis of L1-related INs with substantial axon extended within L1. These comprise L1 INs with soma, dendrite and axons in L1 (left); L2 and L3 INs with axon into L1, and the “Martinotti” cells (Nigro, Hashikawa-Yamasaki et al. 2018, Wu, Sevier et al. 2023) from L5 (right). **(H)** Output of these L1-related IN types onto ExN types, reported for synapses made in L1. Note differential target profiles, also for the subtypes of Martinotti neurons. **(I)** Is the L5-related and L1-related axonal targeting by Martinotti- type neurons coordinated: Are the same neurons targeted systematically? Connectomes shown for output synapses of Martinotti INs made within L1 and those made below L4, for same population of putative postsynaptic ExNs. Hadamard product (right) reports coordinated innervation by individual Martinotti axons. ExNs receiving coordinated innervation in L1 and L5 (row sums of product matrix) were slightly more prevalent than in randomly shuffled connectomes for L5B targets, but not L5A (bottom; number of coordinated innervations per Martinotti neuron rank sorted; rand. Refers to same analysis after random permutation of IN output in the L5 innervation matrix). **(J)** Examples of coordinated innervation of L5B ExNs by individual Martinotti INs (two IN synapse sets shown per target, blue, red, respectively). Even in coordinated innervation cases, the number of involved synapses is small (3-5), suggesting localized dendritic effects rather than global inhibitory impact onto the postsynaptic neuron. **(K)** Who is driving Martinotti INs? Input from ExN types. Note strong innervation by L2HC pyramidals of one Martinotti subtype, inset shows example of this connection across many cortical layers (L2->L5 IN). **(L)** Input to Martinotti cells from INs.

### Layer 1 inhibitory interactions

In addition to long-range axonal inputs, L1 is innervated by inhibitory axons both from INs within L1 and those from deeper layers with substantial L1 projection (note that excitatory axons typically do not reach L1 within the column). Fig. 5G,H report those IN types that have substantial axonal projections in L1, and their output to ExN types made by synapses within L1 (i.e. excluding the outputs made in deeper layers, if any, see below). ExN targeting showed a certain level of specificity, in particular among the population of likely Martinotti INs, of which one had a preference for L2 innervation, while the other for L3+L5B innervation (Fig. 5H).

We next asked whether the fact that Martinotti INs have axonal outputs both in deeper layers *and* in L1 was used for a coordinated innervation of L5 pyramidal neurons, which also have dendrites both in deeper layers *and* L1 (Fig. 5I,J). For this, we restricted the output connectome of Martinotti INs to synapses made below L4, and to synapses made in L1 (Fig. 5I, left) and binarized these two connectomes. We then element-wise multiplied these two connectomes, yielding a matrix reporting for each presynaptic Martinotti axon and each postsynaptic ExN whether the ExN received coordinated innervation by that IN in lower layers *and* L1 (Fig. 5I, right). The row sums of this matrix yielded the number of ExNs receiving coordinated innervation from a given IN. These were then rank sorted and compared to random shufflings of the lower-layer innervation matrices (randomly permuted along rows). We found that for L5B ExNs, the actual innervation matrices showed a higher level of coordinated innervation than random comparison, but not for L5A ExNs (Fig. 5I, bottom right). We then looked at examples of such coordinated innervation of L5B ExNs by Martinotti INs (Fig. 5J). Finally, we asked about the ExN and IN inputs to Martinotti-type INs within the column (Fig. 5K,L), finding differential activation from ExN types (Fig. 5K) and mostly mutual inhibition among Martinotti INs (Fig. 5L).

### Main signal flow within the cortical column

Based on these analyses, we were now interested in the main excitatory and inhibitory signal flow within the cortical column (Fig. 6). First, we computed the type-to-type connectivity matrices for E-E (Fig. 6A), E-I (Fig. 6B), I-E (Fig. 6C), and I-I (Fig. 6D) reporting the mean number of synapses made by all neurons of a given presynaptic type onto a single neuron of a given postsynaptic type. As we had seen before (Fig. 4E,L), this measure promises to report the biophysically plausible effect a given presynaptic population can have on the neurons of a postsynaptic population. Other possible and previously used measures, such as connectivity rate or unitary connection strength, cannot as directly be interpreted as reporting the ability of a presynaptic population to excite or substantially inhibit a postsynaptic population. In addition, this measure can be additively combined for a given postsynaptic neuron type: if, for example, one doubts the subdivision of L3 ExNs into L3A,L3B,L3C, one can simply add these inputs for a given postsynaptic type and obtain a valid readout of the combined effect. These matrices report main type-to-type connectivity, they may underreport the effect of sparse presynaptic populations. The inputs from VPM and L1 were also added to the matrices in Fig. 6A,B (-> ExN, -> IN, respectively).

**Figure 6:**
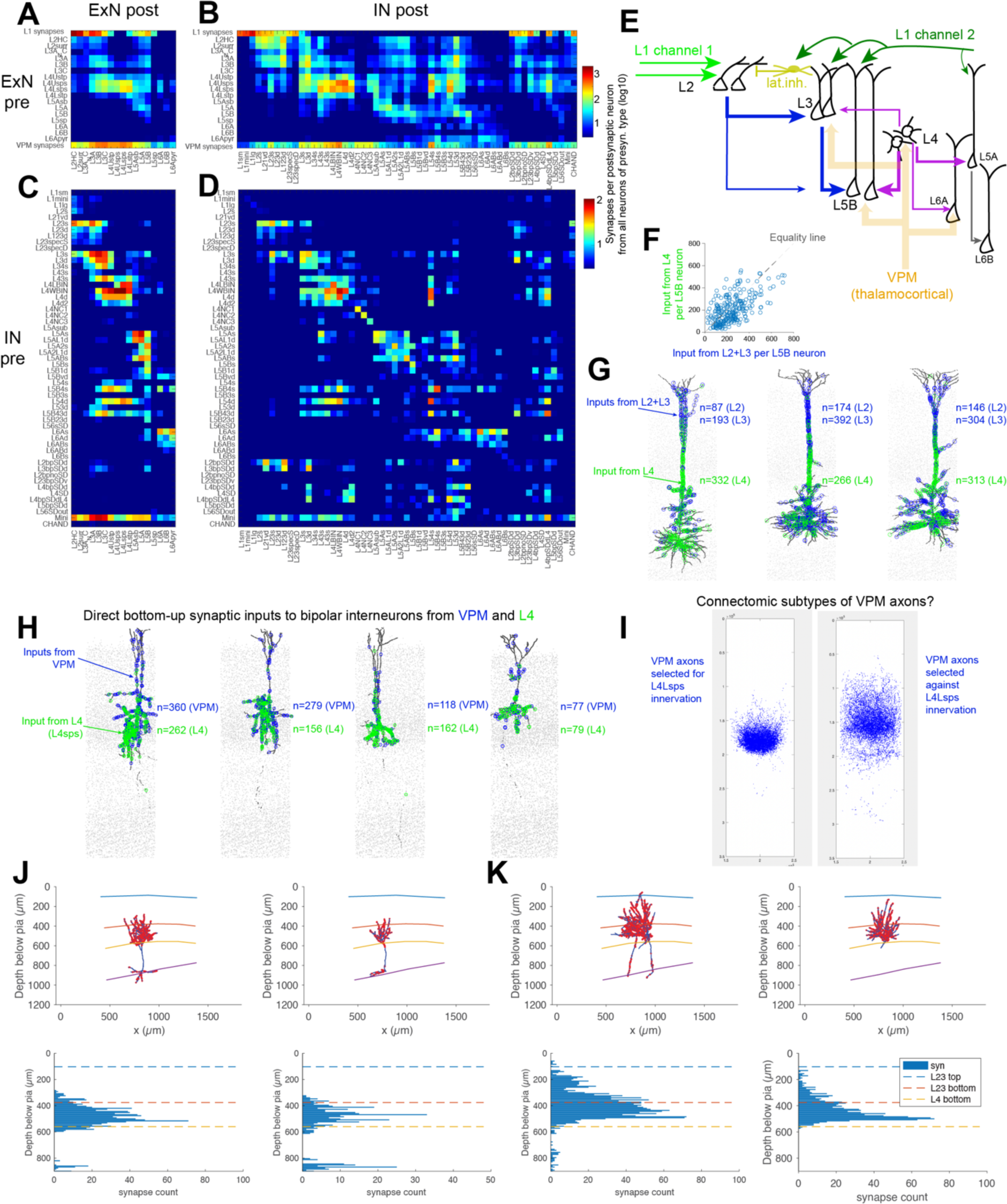
Excitatory-inhibitory signal flow in the cortical column. (A-D) Cell-type based connection matrices for E-E (A), E-I (B), I-E (C) and I-I (D) connectivity. Note connection strength is reported as average number of synapses per postsynaptic neuron made by all neurons of a given presynaptic type, a readily interpretable measure of the effect that a presynaptic neuronal population can possibly have onto the postsynaptic neurons. If this number is small (i.e. below reasonable physiological effect), it is unlikely that the presynaptic cell type can have systematic and reliable effect on the postsynaptic cell type. VPM input (bottom row in A,B) and L1 input (top row in A,B) also reported. Numbers are lower bounds due to reduced axon recall; note that numbers can be added for multiple presynaptic types onto a given postsynaptic type for probing of neuron type recombinations. **(E)** Resulting main signal flow in the cortical column. Note first the separate L1 input channels, likely coupled by lateral inhibition (see Fig. 5 analyses) and clear separation of L2 from L3 by differential circuitry (see A-D). Note further that overall VPM activation is substantial in all layers except L2. Furthermore, dedicated L4->toL6A and L5A->toL6B circuits are found (see A). Finally, the input to L5B neurons (considered a main output of a cortical column) is as strong from L3 as it is from L4. Here, the fact that L4 has many more neurons, even if weaker individual connections, is relevant and yields similar convergent synapse numbers per postsynaptic neuron. The combined effect of presynaptic neuronal prevalence and per-connection synapse number can be clearly seen **(F)**. **(G)** Examples of L5B pyramidal neurons and all their L2, L3 and L4 input synapses, showing the extent to which L4 directly innervates L5B pyramidal neurons, short-circuiting the previously assumed serial canonical cortical circuit (L4->L3->L5B). High recurrence at all levels is found. **(H)** Does the assumed distinction in bottom-up and top-down driven IN populations hold? Strong direct innervation of bipolar INs by VPM input. These INs were thought to be driven primarily by top-down inputs. However, they are activated by VPM directly, by L4, and other processing layers, revoking a clear differentiation into bottom-up and top-down driven interneurons. See Fig. 4 for direct effect on columnar signal gating by activation of these bp-INs. **(I-K)** Identification of separate VPM input channels. First, automatically reconstructed VPM axons were separated by their propensity to innervate L4Lsps. These showed differential axonal spread. To control for selection bias, VPM axons were manually reconstructed at whole-axon level. Whole-axon reconstruction examples reveals two subtypes of VPM input axons: one clearly restricted to L4, and another with strong innervation in the upper L4 and L3+2. This distinction may be related to centrally located VPM neurons (Zhang, Wang et al. 2021). This could represent two separate VPM input channels into the column.

Based on these matrices and the previous analyses in Figs. 1-5, the main signal flow is summarized in Fig. 6E. L1 input can be considered as at least two channels, one with preference for input to L2 ExNs, one with preference for L3+L5. The differential innervation of L1lg and L1mini INs by the non-L2 preferring L1 axons (Fig. 5F) together with these IN types’ preference for L2 ExN inhibition (Fig. 5H) can be interpreted as a directed lateral inhibition circuit between the two L1 input channels (Fig. 6E, top). The L2 signal is propagated downwards into the column primarily to L3 ExNs and to L5B ExNs. VPM input, when considered as one main input to the column, innervates most ExN types, but less so L2 ExNs. L6 ExNs are differentially innervated by L4 for L6A and by L5A for L6B ExNs (see Fig. 6A and 6E).

When analyzing the input to L5B ExNs, considered one main output channel of the column, we found that quantitatively, the input from L3->L5B is of similar magnitude as the input directly from L4 (Fig. 6F,G). This is in stark contrast to the canonical circuit model, in which L5B output is primarily driven by L2+L3 inputs, and is evidence for a powerful shortcut in the intracolumnar circuit. Fig. 6G illustrates the strength of the L4->L5B connection. This can be considered another example for the fact that unitary connection strength alone (which is smaller for L4->L5B) is not sufficient to judge main signal streams; rather, the prevalence of the presynaptic populations is of critical importance.

We also found that contrary to the standard circuit picture of inhibition and disinhibition, bipolar (bp) INs with strong disinhibitory outputs (likely VIP-like INs), are directly and strongly innervated by VPM and L4, thus the earliest bottom-up stages of cortical processing (Fig. 6H). While we did find evidence for some bipolar subtypes receiving less intracortical activation (Fig. 4M), this direct innervation of bipolar INs by bottom-up inputs questions the sequential IN recruitment picture that has emerged from the circuit studies over the recent decades (e.g. (Yu, Hu et al. 2019)), and may explain their functionally selective responses (Guy, Mock et al. 2023).

Finally, we wondered whether the VPM input axons are a homogeneous population themselves. First, we selected VPM axons by their output preference for/against L4Lsps innervation (Fig. 6I) and found a possible split into L4L-focused vs. upper layer-focused VPM axons. This could however be caused by splits in VPM axons, yielding local biases depending on the chosen target selections. To test whether a subdivision of VPM axons is plausible, we manually reconstructed VPM axons to full extent in the column (Fig. 6J,K), carefully distinguishing real from truncated endings, and found that indeed, two subpopulations could be readily identified with one focused on L4 (Fig. 6J), and one extending to upper layers (Fig. 6K) while avoiding synapses in the bottom third of L4. This subdivision could relate to VPM center vs. margin neurons (Zhang, Wang et al. 2021), and it may indicate the need for a more detailed study of VPM specificity within the column. For example, a L4 focused VPM channel would not recruit as broadly as the overall VPM population, thereby allowing a dedicated signal flow into the column with higher target specificity.

### Top-down *vs* bottom-up circuitry in the column

A hallmark of presumed cortical processing is the combination of sensory input with pre-assumed hypotheses or predictions. For a given cortical area, this is assumed to be represented by bottom-up input (mainly to layers 4 and 6), and top-down input (mainly to L1). We next wanted to investigate how this assumed convergence of data streams could be implemented at the cortical column level (Fig. 7). First, we considered again the detailed ExN-ExN type connectome (Fig. 7A). We also summarized the number of neurons in each ExN and IN type (Fig. 7B,C). When considering the representations in the cortex as spatial codes within homotypic populations of neurons, the number of neurons in a relevant cortical module can be considered an upper bound on the representational capacity of each neuronal population. Obviously, ExN-based representations offer far larger capacity than IN-based representations, which would, if used for encoding, constitute substantial representational bottlenecks (Fig. 7C).

**Figure 7:**
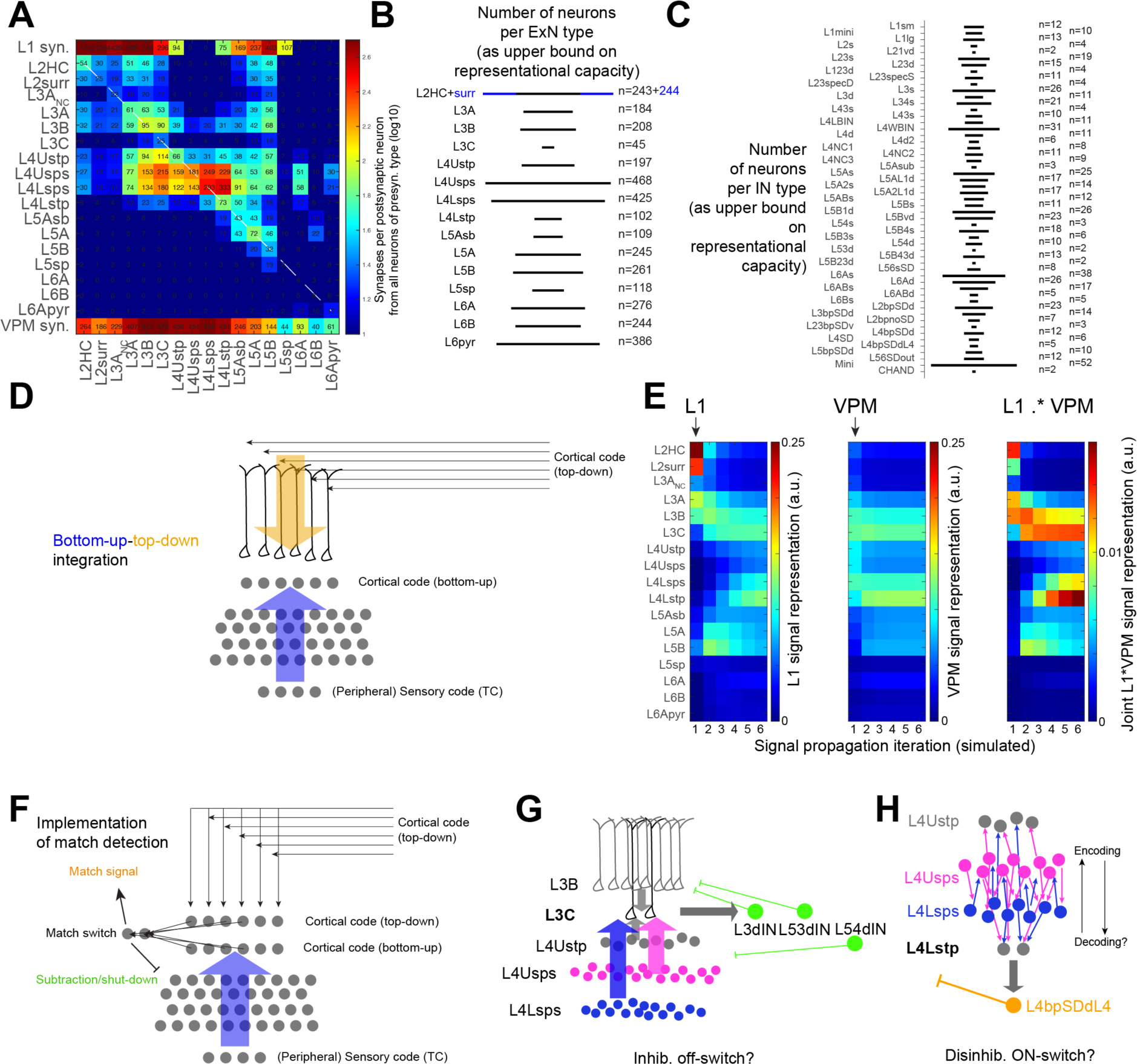
Possible encoding and switch circuits, representational capacity of ExN and IN types, and summary of predicted information flow. **(A)** Detailed ExN- ExN matrix (same as in Fig. 6A) with mean number of synapses onto postsynaptic neuron reported. **(B,C)** Number of neurons per neuron type, the upper bound of representational capacity per neuron type. Note that except for L2,4 and 6, cell type capacity is about 100-250 per column, indicating a maximum dimensionality for a possible cortical code. Note that INs (right) are mostly very rare, creating representational bottlenecks, in particular for bipolar -type INs. **(D)** Sketch of the possible integration of bottom-up and top-down information in the column. **(E)** Simple stepwise iterated matrix multiplication (normalized after each step) of L1 input (left) iteratively multiplied to the ExN-ExN type connectome; and VPM input (middle) to the columnar network. Rightmost matrix reports Hadamard product of L1 and VPM flow matrices. Note main convergence after 2-3 iteration steps in L3B/C, L4stp and L5B (intermittently), positioning these populations as ideal combination points for these information streams. **(F)** If one considers the matching of intracortical (predictive) encodings with sensory-evoked current data as a goal of columnar circuitry, representational matches need to be computed. This can be either done directly in the apical tufts vs. basal inputs of pyramidal neurons (see D,E) – or via switch circuits, in which a small number of neurons is densely innervated by the comparative populations, and gets active once a sufficient number of matching inputs is received. The output of the switch population would be interpreted either as a match signal, or trigger a subtractive (inhibitory) signal that subtracts or shuts down the matching (predicted) input. **(G,H)** Search for low-N possible switch populations: a possible OFF switch via L3C (G) and a possible ON switch via disinhibitory circuits for the possible matching of encoded and decoded peripheral sensory representations (H). L5sb is an additional possible switch population.

How would the top-down inputs be able to converge with bottom-up projections? One view of such convergence is that the pyramidal cells themselves, in particular those with access to L1 *and* VPM innervation, could integrate these two inputs at the single-cell level, using mechanisms for coincidence detection (Fig. 7D, (Larkum, Zhu et al. 1999, Larkum and Zhu 2002, Waters, Larkum et al. 2003, Larkum, Senn et al. 2004, Larkum 2013, Major, Larkum et al. 2013)).

Alternatively, the inputs could gradually be transmitted within the column, finding possible sites of convergent recombination. We first used a simple simulation of signal propagation using the input strength from L1 to all ExN types and the input strength from VPM to all ExN types (Fig. 7E), and multiplied this iteratively to the ExN-ExN connectome. As expected, L1 input is strongest in L2, and then propagates to L3 an L5, followed by possible propagation to L4stp. VPM input is more ubiquitous. For explicitly considering combined L1 + VPM representations in this simplified picture, we elementwise multiplied the two matrices (Fig. 7E right), yielding in particular L3C ExNs and L4Lstp ExNs as possible sites of convergence after 4 signal propagation iterations. These are also the sparsest ExN populations.

When explicitly considering the possibility of an intracolumnar matching of top-down and bottom-up cortical representations (Fig. 7F) via circuit-level implementation, a plausible solution is the occurrence of switch populations which detect the possible match of sensory input and top-down prediction (Fig. 7F). Such switch populations are ideally quite small in number, such that even sparse presynaptic representations can converge to excite sufficient amounts of switch neurons. Then, the match signal can either be propagated itself, or be used as a trigger for inhibitory (subtractive) circuitry according to the concept of predictive coding (Friston and Kiebel 2009, Keller, Bonhoeffer et al. 2012, Leinweber, Ward et al. 2017, Keller and Mrsic-Flogel 2018, Keller, Dipoppa et al. 2020, Schuman, Dellal et al. 2021).

In fact, when searching for sparse switch populations, L3C and L4Lstp are noteworthy. When analyzing the input and output circuitry of each (Fig. 7G,H), we found that L3C is strongly innervated by L4 populations, making it plausible to obtain the L4- transformed bottom-up representations; and also receive sufficient top-down input both directly and indirectly (see Fig. 7E). The output of L3C to certain types of INs is strong enough to plausibly trigger their activation, which by the INs’ target properties would plausibly shut down or inhibit the L4 and L3 excitatory networks (Fig. 7G). This could be interpreted as a possible inhibitory subtraction, along the logic of subtracting/inhibiting a match between prediction and sensory input. For the L4Lstp population, conversely (Fig. 7H), a key output is a disinhibitory IN type that could possibly be interpreted as allowing a match signal to pass. In particular since the input to L4Lstp is primarily from L4 in the early steps of input to cortex, and the intra-L4 networks have the resemblance of an encoding circuit to transform VPM input representation into a possible cortical code, one could speculate this switch signal to report an encoding success rather than a prediction match signal.

### Detailed cellular-level structure of the L1 top-down projections

In the preceding analyses, signal flow within the column has been considered at cell-type level. However, to implement a possible direct prediction from higher-order cortical areas to the sensory cortex, high specificity of connections has to be maintained. We next analyzed whether evidence for such cellular-level specificity can be found (Fig. 8).

**Figure 8:**
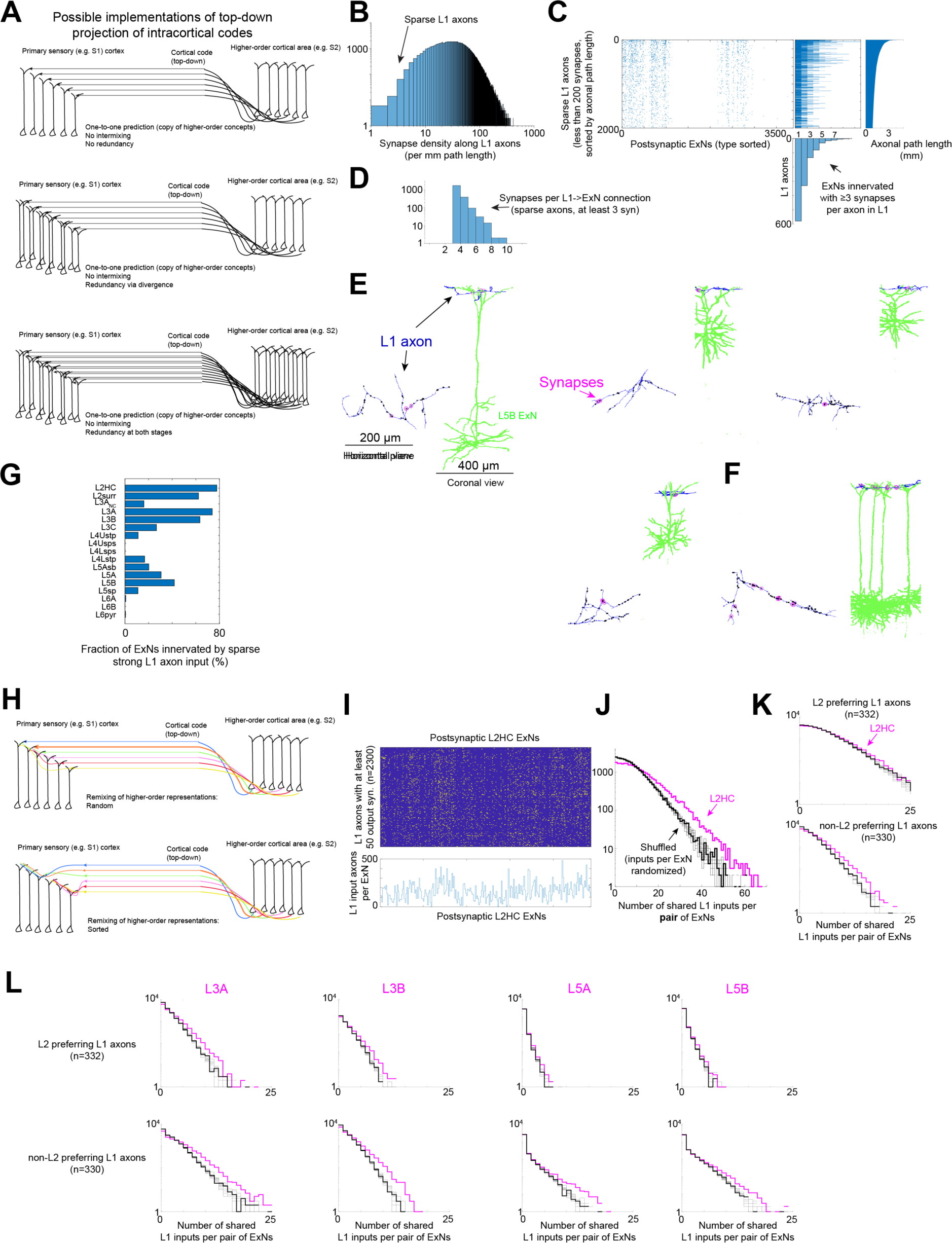
Detailed analysis of possible top-down information flow via L1 into the column. **(A)** Possible circuits for projecting a higher-order generated prediction into the lower-order sensory cortex; these models assume that the encoding in the higher cortical area is similar to the one in the lower area, thus one-to-one matching of innervation would be appropriate. This predicts very sparse innervation within L1: axons that may target only one or a few intracolumnar neurons but ideally with strong connections. Three variants of redundancy shown. **(B)** Search for sparse but strong connections within L1. **(C)** Connectome from 2100 L1 axons (sorted by path length to identify sparse axons) to all ExNs, only connections with at least 3 synapses reported. Right, Number of strongly innervated ExNs per L1 axon, note in fact predominance of 1-target axons. Note however that axons make 1- and 2-synapse connections more promiscuously. **(D)** Distribution of strength of connections (at least 3 synapses) shows that connections can be strong but are mostly of intermediate synapse number (3-5). **(E)** Examples of L1 axons targeting one ExN in the column with at least 3 synapses. **(F)** Example of L1 axon targeting n=4 L5A ExNs via at least 3 synapses each, indicating possible redundancy. **(G)** Are ExN populations systematically targeted by the sparsely innervating L1 axons? For this, we analyzed the fraction of ExNs of a given type that received sparse but strong (at least 3 synapses) L1 innervations. Note very strong coverage for L2, L3A+B (but not L3C) ExNs, and less strongly for L5 pyramidal neurons. **(H)** Alternatively, projections from higher cortical areas could intermix onto the columnar pyramidal neurons, creating novel projections in coding space; this could be systematic (bottom) or random (top). **(I)** Analysis of joint innervation of pairs of ExNs by multiplets of L1 axons. **(J)** Distribution of number of shared L1 innervations over pairs of L2HC, compared to a random shuffling of the connectome in (I) along the columns (i.e. input shuffling). Note higher prevalence of jointly innervated ExN pairs compared to random. **(K)** When analyzing the previously identified L1 axon subtypes separately, however, no additional systematic shared innervation was found. **(L)** Detectable but weak shared innervation bias for axons in L1 to various ExN types. Together, a direct projection mapping is more likely than remixing, or a combination depending on L1 axon type.

If there exists a common cortical code, in which populations of excitatory neurons in higher cortical areas encode concepts that are transformed into sensory-cortex-level predictions and projected one-to-one to matching populations of excitatory neurons in lower cortical areas, the input within L1 should be very sparse (Fig. 8A), even when considering various levels of redundancy. Conversely, if one assumes higher-order representations to be projected into the lower-sensory-cortex encoding space by intermixing (Fig. 8Hff), different projection types are expected. We looked for evidence of both of these concepts, which could of course also be implemented in parallel.

First, we specifically looked for axons within L1 that had substantial path length (1mm and more) but few synapses – a direct prediction from the one-to-one top-down encoding model described above (Fig. 8B,C). We then asked which ExNs were innervated by such axons, and whether any of these innervations had a sufficient number of synapses per connection, such that a direct triggering of distal dendritic nonlinearities would at least be plausible. For this, we restricted the analysis to connections with at least 3 synapses (Fig. 8B-G). In fact, most sparse L1 axons made only one strong output connection to ExNs (Fig. 8C). A prerequisite for sparse top-down projections of the above nature would however be that while one given axon makes few strong connections, the entirety of such sparse axons innervates entire populations of ExNs in the column. We therefore asked when combining the inputs from many sparse L1 axons onto ExNs (Fig. 8E), what fraction of postsynaptic ExN populations would be innervated (Fig. 8G). We found that up to 80% innervation could be reached for L2 and L3 ExN types. We also found examples of subtype-specific innervation in L1 (Fig. 6F) with sparse L1 axons making strong outputs to 4 L5A pyramidal neurons, a pattern seen in several cases. For further analysis of such redundant innervation, L1 axon merge rates will be further reduced.

Finally, we analyzed the alternative implementation of long-range projections intermixing onto columnar ExNs (Fig. 8H-L). Here, we distinguished the case of coordinated mixtures from random projections (Fig. 8H). In the case of coordinated input mixing, one would expect sets of ExNs to share a number of L1 innervations, while for random intermixing such coordination would be absent. To analyze this, we used L1 axons with at least 50 outputs, and investigated for each ExN type whether coordinated above-random innervation in L1 was found (Fig. 8I-L). As a fist measure, we analyzed the pairwise input correlation between pairs of ExNs (Fig. 8J) and compared this to randomly shuffled versions of the connectomes. While the overall L1 input to L2HC ExNs showed some above-random coordinated input (Fig. 8J), this was fully accounted for when using the L1 axon subtypes defined earlier (Fig. 8K; see Fig. 5C,D). Also for other ExN types, no strong evidence for coordinated L1 innervation could be found (Fig. 8L).

Together this may indicate that indeed, a subset of sparse L1 axons provides very specific input to ExN populations, possibly along the lines of a top-down projection of higher-order cortical encodings.

### Higher-order structure within homotypic ExN connectomes

We next wanted to study whether also at the level of intracolumnar ExN networks, structured connectivity can be found within cell-type homotypic connectomes. One possible structure within homotypic networks are subclusters. These were identified in the initial cluster analyses for ExN type definition, and some additional spatial structure was found.

However, a key question is whether other types of structured connectivity can be detected, in particular feedforward network structure, either as overall feedforwardness or band structure indicating more specific synaptic chains.

To test for this, we applied an efficient sorting algorithm recently published by Borst (Borst 2024). With this, we could efficiently search for sortings of ExN-ExN connectomes that would yield low recurrence (i.e. high feed-forwardness). When applying this to ExN connectomes within layers (Fig. 9A-C), we indeed found lower degrees of recurrence in optimized sortings compared to random Erdös-Rényi (ER) networks (Fig. 9E). The obtained sortings were not obviously related to spatial arrangements (Fig. 9D), making a pure margin effect unlikely. Yet, random degree-preserving reshuffling of these matrices also yielded low-recurrence sortings (Fig. 9H). We therefore had to test whether 1) the obtained sortings had any additional biologically relevant implications and 2) whether the obtained sortings had evidence for feedforward bias even for medium-out-degree neurons in the middle of the sorting sequence. When we analyzed the translaminar connectomes of L4 and L3 populations, we indeed found that the feed-forward sorting in L4Usps induced an inverse degree of connectivity to L3A (Fig. 9F,G). This was counterintuitive, since neurons early in the sorting would generally be biased for higher out degrees than those later in the sorting; yet, the connectivity to L3A was inverted (low early in the sorting, higher later), yielding >2-fold difference in L3A innervation. This could be interpreted as a biological support for the otherwise statistical sorting, even if a large number of possible effects in the translaminar connectome were investigated.

**Figure 9:**
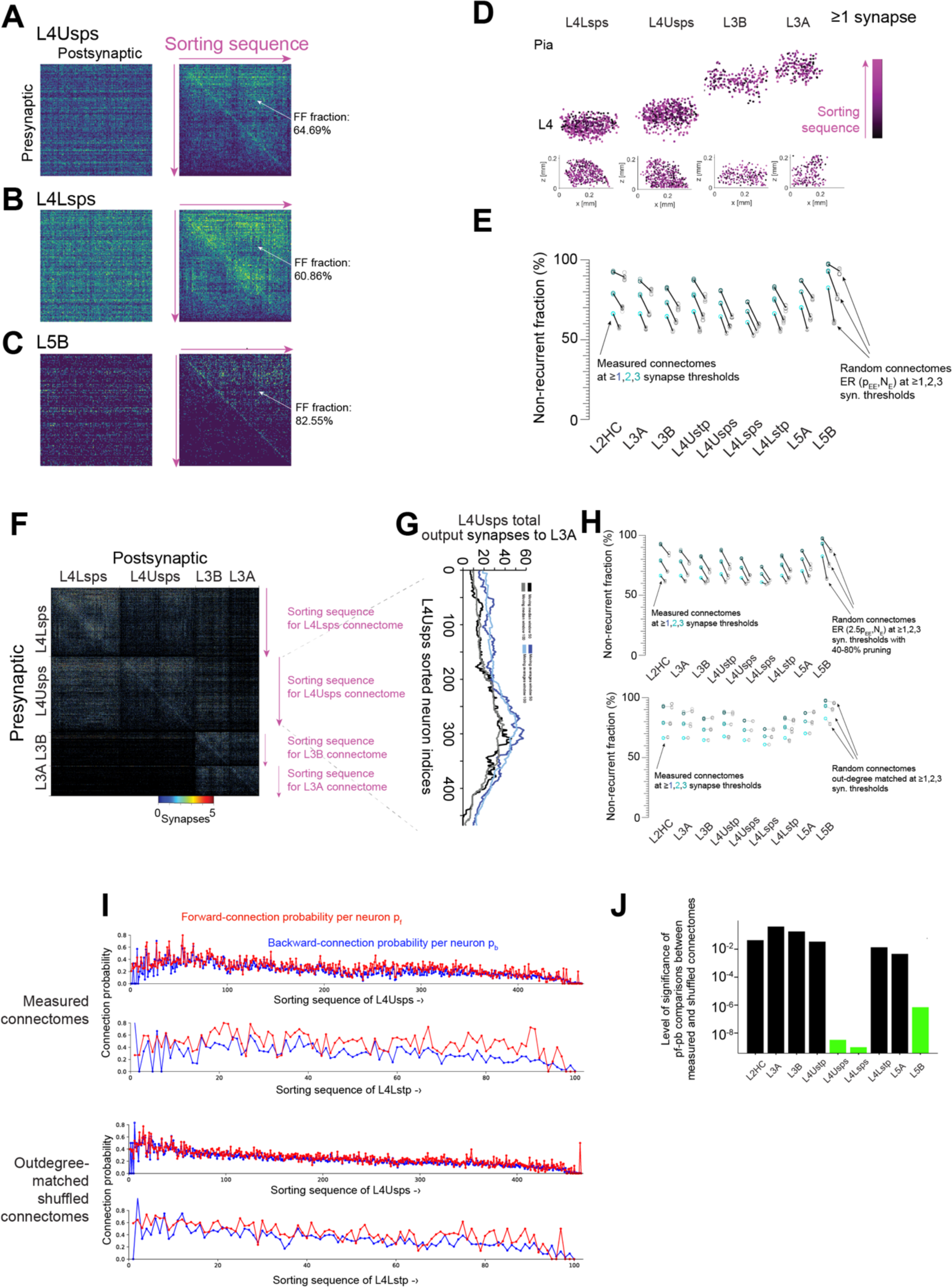
Analysis of substructure within homotypic E-E connectomes. **(A-C)** Search for feedforward sorting within EE connectomes using a fast sorting method (Borst 2024). Feedforwardness (defined as 1-recurrent fraction) of connectomes after sorting was 60-80% (A-C). **(D)** Sorting could not be explained by simple spatial (e.g. margin) effects. **(E)** Feedforwardness was higher than that for random Erdös- Renyi connectomes (ER) with matched pairwise connectivity. **(F,G)** Investigation of translaminar connectivity between L4 and L3, with each ExN connectome sorted by feedforwardness within each subtype. Note noticeable gradient of innervation from L4Usps to L3A that was antiparallel to the feedforward sorting: the later in the sorting chain for L4Usps, the stronger the connection to L3A (overall a more than two-fold change, G). This may indicate that the weak feedforwardness in these networks is used for input-output related connectivity biases. **(H)** Controls: possible effect of axonal pruning on apparent feedforwardness (top). When pruning a random ER network by 40-80%, the resulting feedforwadness is not larger than the underlying width of the outdegree distribution, still yielding less forwardness than the measured connectomes (top). Out-degree matched random connectomes, however, are largely similar in feedforwardness (bottom). **(I,J)** We therefore asked whether the probability of forward connectivity pf is larger than the probability of backward connectivity pb also for neurons placed in the middle part of the sorting (where extreme differences are unlikely). I shows pf and pb for L4Usps and L4Lstp (top) and the outdegree- matched shuffled connectomes. For L4Usps, L4Lsps and L5B, the differences are strongly significant (J), indicating an actual feedforward sorting beyond outdegree- variability matched random controls.

One concern was that the sorting was induced by effects of axonal pruning, creating artificially broadened outdegree distributions. However, in a ground truth comparison tracing, we found a variability in the number of output synapses of 50-70%. When testing the effect of axonal pruning on such degree distributions, the feedforwardness would not be induced by pruning, as long as the variability of axonal pruning would not exceed the underlying degree variability (Fig. 9H).

Finally, for the statistical test of forward-bias even for neurons with average outdegree, we found that indeed, the probability of forward connections exceeded the probability of backward connections in the sorting, also for the least biased members of the sorting (Fig. 9I,J).

Together this may indicate that a certain degree of feedforward structure can be found in intracortical ExN connectomes, in particular for L4Lsps and L4Usps and L5B, which is primarily driven by outdegree variability, but may be exploited for biased translaminar connectivity.

### Traces of Hebbian plasticity in the column

As a final set of analyses, we wanted to investigate whether the traces of homosynaptic plasticity (in particular, Hebbian LTP), which we had previously extracted in a 1000-fold smaller connectomic reconstruction (Motta, Berning et al. 2019), using paired-synapse analysis concepts from (Bartol, Bromer et al. 2015), would be present in other connections within and into the column, as well (Figs. 10,11). We first analyzed the overall (Fig. 10A) and connection-specific (Fig. 10B) distributions of synapse sizes. Input synapse size distributions were more homogeneous than output size distributions, possibly representing an input but not output normalization effect. When explicitly analyzing the correlation between average input and output synaptic size across cell types (Fig. 10C), we found much more narrow input than output size distributions in all types except L5 and L6 ExNs.

**Figure 10:**
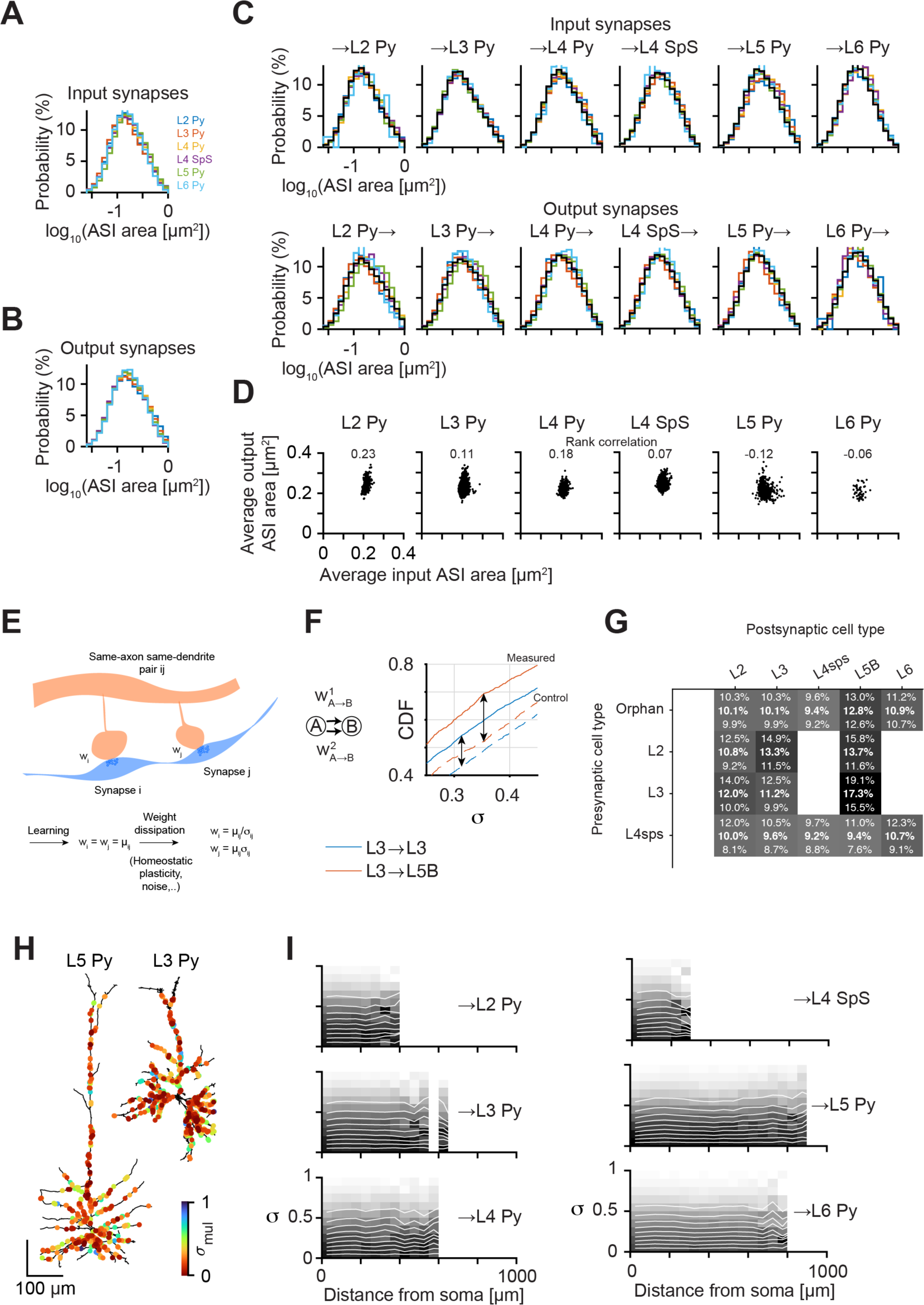
Analysis of synapse sizes (strengths) and traces of Hebbian learning for intracolumnar connections. **(A,B)** Overall distribution of input and output synaptic sizes. Synapse size distributions per connection. Note that inputs onto a given cell types follow similar size distribution, irrespective of source. The outputs of a given cell type, however, depend significantly on the postsynaptic target. **(C)** Detailed comparison of source- or target-dependent synaptic size distributions. Input synapse sizes are more identical than output (may be more strictly controlled/normalized). **(D)** Relation between input and output synapse size (average ASI) across ExN types shows narrow input but broader output size distributions, except for L5 pyramidal neurons. **(E)** Analysis of same-axon same-dendrite synaptic pairs and their synaptic weight: assuming a shared (but unknown) homosynaptic target weight, perturbed by processes acting on each synapse separately. See methods for detailed derivation. **(F)** Example comparison of the same-axon same-dendrite vs shuffled synaptic pairs (measured vs control). Arrows indicate the maximum difference between the cumulative distribution function, which corresponds to the surplus of low-noise connections. **(G)** Analysis of overrepresented low-perturbation synaptic connections for various connection types in the column (only those with significant over-similarity shown). Numbers report most likely and 5^th^..95^th^ percentile estimates for the fraction of synaptic configurations that are overabundantly similar (a more conservative measure for learnedness than the one reported in (Motta, Berning et al. 2019)). Note almost 2-fold difference between weakest and strongest learning effect: by far the strongest is the L3-to-L5B connection, almost 2-fold more similar than L4sps-to-L5B (the two main drivers of L5B, see Fig. 6E-G). Note further that also the external (“orphan”) connections are most strongly synaptic weight-homogenized for connections to L5B. **(H,I)** Location dependence of synpatic size similarity in joitn same- axon same-dendrite synaptic pairs. No obvious distance dependence found except a weak effect for L4 spiny stellate neurons.

**Figure 11:**
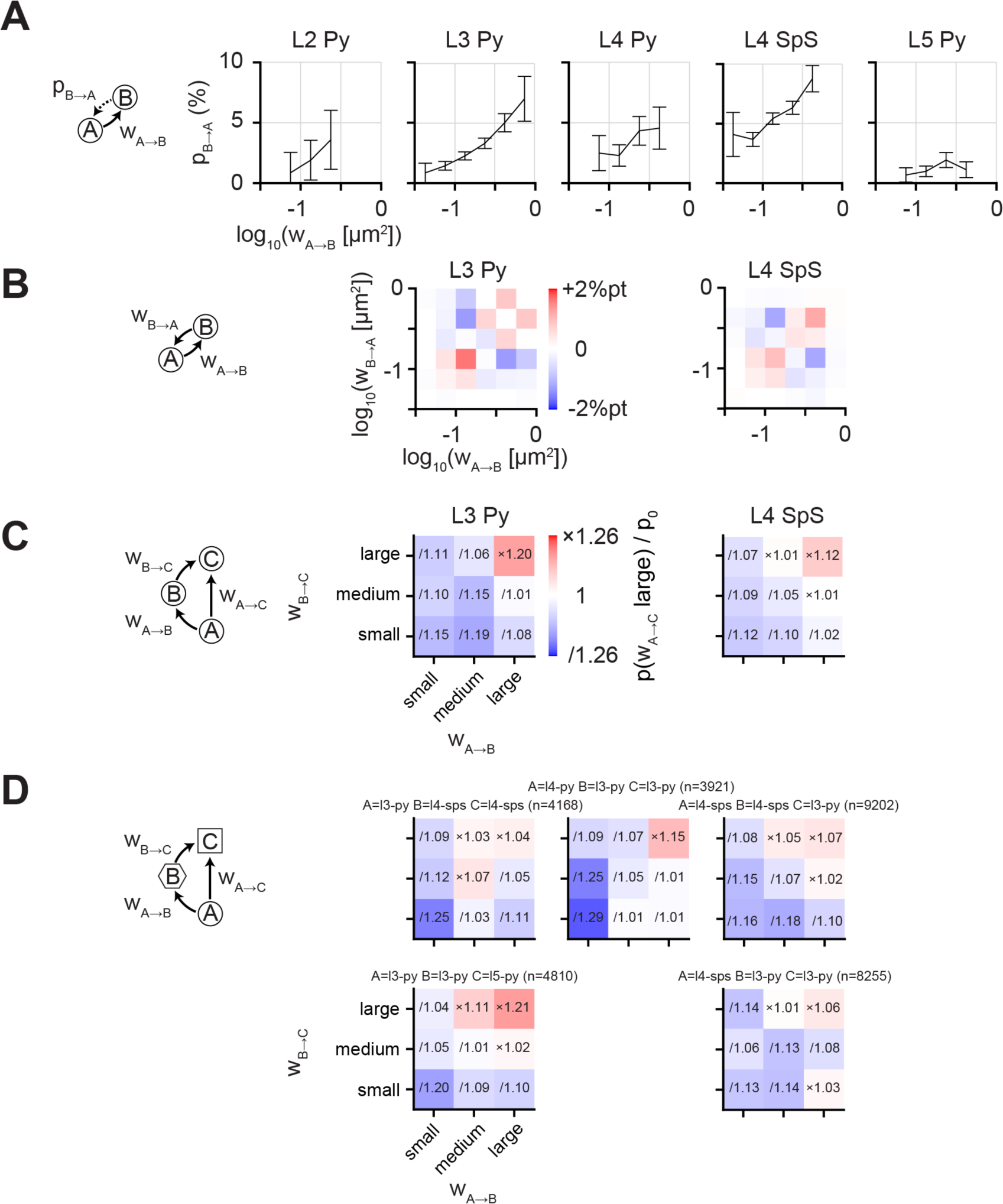
Analysis of homosynaptic plasticity in reciprocal and triplet connections. **(A)** Probability of reciprocal connection in dependence of forward synaptic size. Note strong correlation for all connections except L5 pyramidal cells. **(B)** Pairwise recurrent connections have aligned synaptic weights, in conflict with timing- dependent Hebbian plasticity, and in favor of correlation-based LTP, analyzed for L3ExNs and L4sps. **(C,D)** Analysis of possible transitivity predicted by correlation- based learning in synaptic forward triads. Large synaptic size in A->B and B->C is correlated to large (strong) connections in A->C.

Then, we used an improved model of detecting evidence for homosynaptic plasticity (Fig. 10D) in which it is assumed that pairs of same axon-same dendrite innervations converge onto a joint synaptic weight (i.e they share a common synaptic target size), but get perturbed by various sources of multiplicative noise (e.g. homeostatic (but heterosynaptic) plasticity), yielding states of underlying target weight *µ* and synaptic noise magnitude *σ*. We then obtain distributions of the maximum likelihood estimates of the target weight and noise magnitude for each set of connections (see Methods), and compare it to randomly shuffled weight pairs (Fig. 10E). With this, we obtained lower-bound estimates of the degree of homeosynaptic weight convergence (e.g. Hebbian plasticity) over the various connections in the column (Fig. 10F; note these estimates correspond to the above-random portion of the weight variability histogram, see Fig. 7H in (Motta, Berning et al. 2019)). Notably, the amount of surplus low-noise connections varies almost two-fold across connection types (9.2% in L4sps-L4sps connections vs 17.3% in L3->L5B). The connection from L3 to L5B, one of the main output lines from the column, is the most consistent with homosynaptic plasticity, followed by the L2->L5B connection. We also investigated the connections made by axons not connected to a soma in the volume (“orphan” axons), which contain long-range inputs to the column (but are of course confounded by more proximal but pruned source neurons) and could be a primary site of large-scale information storage (Shan 2023). For these connections we found average levels of “learning”, with again L5B neurons receiving the most strongly over-similar inputs.

We then wondered whether synaptic pairs would vary in their weight similarity along the dendrites of given ExNs (Fig. 10G,H), but found no strong relation to distance from soma, except for a modest distance effect in L4sps neurons (Fig. 10H).

Finally, we wanted to make use of the size of the connectome to analyze higher-order synaptic configurations for their possible synaptic weight homogeneity (Fig. 11). In particular, we were interested in whether reciprocal connections had similar or anticorrelated weight targets; and how synaptic triads would behave. We wanted to test the predictions of a particular model of homosynaptic plasticity, namely the covariance rule (or correlation-based learning) in (Sejnowski 1977). In contrast to most other models of homosynaptic plasticity (e.g., spike timing-dependent plasticity), the covariance rule is symmetric. That is, the predicted weight changes for reciprocal connections (A->B and B->A) are identical.

First, we analyzed the reciprocal connectivity in pairs of homotypic ExN connections in dependence of the size of the forward connection (Fig. 11A). We wanted to analyze whether the probability of a connection being reciprocated depends on its homosynaptic target weight (µ). We found a strong dependence in all but L5 connections. Then, we analyzed the relation between forward and backward target weights for reciprocal connections between L3 and L4 ExNs (Fig. 10B) and found strong co-dependence, consistent with predictions of symmetric models of homosynaptic plasticity.

Then, when analyzing triads of connections, we again found that large weights in two subsequent steps of a chain go along with large weights in the direct connection (Fig. 11C), also for translaminar connections (Fig. 11D).

## DISCUSSION

Based on the connectomic reconstruction of close to 10^4^ neurons and their connections, we were able to 1) define a cortical column as a property of the connectome 2) define cortical cell types by connectomic parameters; 3) determine parallel feedforward inhibitory systems within the column; 4) determine the key disinhibitory pathways, finding strong immediate reccurency; 5) identify the pathways of combination of top-down with bottom-up input streams in the column, thereby providing an input flow model of cortical microcircuitry; 6) determine strong shortcuts to the canonical circuit model, emphasizing immediate recurrency rather than sequential activation in the cortical column; 7) determine structure within homotypic ExN connectomes; 8) determine focus points of homosynaptic (e.g. Hebbian) plasticity, in particular the connections to L5B pyramidal neurons; and 9) obtain evidence that Hebbian synapse consistency supports symmetric (e.g. rate based), not asymmetric (e.g. timing based) plasticity models.

### Column definition

For a first connectome of a cortical column, it was decisive to reconstruct a circuit entity with well-defined boundaries and functional significance. The rodent barrel cortex provides such a unique opportunity of clearly identifiable columnar boundaries in L4, and an interpretation of inputs from the sensory periphery via VPM. Only with such a clear columnar definition, conclusions about representational size of cell networks (Fig. 7), lower bounds on prevalence of rare neuron types (Fig. 3), and column-related circuit structure (Figs. 4-8) could be obtained. While our dataset comprised on the order of 10^4^ neurons, these were distributed across one approx. 60% complete column, three neighboring column-parts, and the intermediate septa. Initially, we considered this a disadvantage of our dataset. However, as can be seen in the cell-type and processing analyses (Figs. 2,3,4), this particular arrangement proved an advantage, since we were able to confirm major cell types in the neighboring columns, thus providing a sampling across n=4 columns, at least for the most significant results.

### Reconstruction accuracy

For this work the automation of reconstruction was essential. With automation, however, the ensuing error rates have to be considered, and analyses have to account for possible confounders based on incomplete or erroneous reconstructions. In our case, even with partial recall of axons (but very high recall of dendrites and spines), the analyses presented here were possible. For example, axonal target specificity, as long as axonal distance-dependence effects are not dominant, can be analyzed with partial axonal reconstructions. Similarly, if the recall rates are sufficiently similar between neurons, comparative convergence analyses can be carried out. Relative impact on postsynaptic neurons can be evaluated. However, the total number of synapses per cell type connection reported here is certainly a lower bound. Similarly, connectivity among ExNs is a lower bound estimate. We have approximated for each analysis to what degree insufficient recall (or in some cases remaining mergers) would affect the analysis results, and only report analyses where this estimate is tolerable.

### Rare interneuron types

One aspect of our analyses is the relative paucity/sparsity of some of the most well-known interneuron types. It had already been indicated in data from visual cortex that for example Chandelier cells were so rare as not be contained in previous unbiased samplings (Schneider-Mizell, Bodor et al. 2021, Schneider-Mizell, Bodor et al. 2024). Here, by sampling at 10^4^ scale, we can determine lower bounds for their prevalence (2 out of 10^4^, thus 0.02%). This is exceptionally rare, and has immediate implications for the functional impact these cell types can have on columnar processing. In particular, if we expect only about 1-3 of these neurons per cortical column, they are almost certain to exert global, not column-precise influence. While in insects and other species with short life spans, single interneurons have been shown to exert outsized influence on the network (Dorkenwald, Matsliah et al. 2023), and thereby the possible demise of a single neuron would seriously alter brain function, such a notion of single-neuron reliability is hard to imagine in the long-living mammalian brain. Here, our data suggests that the required stability of neurons over decades of age is at the level of 1/5000 (i.e. about 3,000 individual neurons for the sparsest type across mouse cortex). Several other proposed cell types (Fig. 3) were of similar sparsity, even if some may be considered part of larger cell type groups.

### Connectomic neuron type definition vs other parameter spaces

We particularly focused on the deduction of cortical cell types from only connectomic data. Such a connectomic-only cell type definition had been successful in the mammalian retina (Helmstaedter, Briggman et al. 2013), where subtypes of a previously lumped-together bipolar type were found by connectomic means, and later morphological differentiation was added (Kim, Greene et al. 2014, Seung and Sümbül 2014). With the quite clear clustering results from the cortical column, we found that at least a large set of cell types can be very clearly obtained by connectomic means, alone. This adds an important cell type definition space, and it will have to be determined whether morphological, electrophysiological and transcriptomic cell type definitions will discover additional subtypes, that connectomic analysis could not reveal. It has to be determined, for example, to what degree long-range target properties of ExNs can be fully predicted by the local connectomic type definitions.

### Vertical extent of columnar structure

While the definition of cortical columns at the level of layer 4 has been undisputed in S1 cortex, the question of the columnar extent into supra- and infragranular layers has remained a matter of debate, with substantial LM-based morphological data indicating that especially lower layers would lose their columnar assignment (in favor, for example, of row- or arc-based alignments, (Narayanan, Egger et al. 2015)). Our connectomic clustering showed, however, that even in the extreme layers (L2 and L6), intracolumnar circuit modules do exist. These are likely driven by the L4-to-L3-to-L2 circuitry (supragranular), and the unexpectedly strong L4-L6A circuitry (infragranular). The only cell type that showed no indication of columnar reference were the sparsely innervated L5 ExNs (L5sp, Fig. 2G).

### Top-down canonical circuit

We were especially interested in the identification of top-down circuits, and found two separate streams in L1 (to L2 vs L3C+L5B ExNs, respectively) and possible sparse one-to-one top-down projections, and in the combination of this input with bottom-up input (Figs. 5-8). Whether this circuitry is similarly applicable to higher-order, non-sensory cortices, will have to be investigated.

### Cortex type

The choice of S1 vibrissal cortex for the first cortical column connectome was not arbitrary: here, as decades of anatomical and functional literature had shown (Welker, Johnson Jr et al. 1964, Woolsey and Van der Loos 1970, Catania and Kaas 1995, Woolsey 1996, Jain, Catania et al. 1998, Lubke, Roth et al. 2003, Helmstaedter, de Kock et al. 2007), the concept of the cortical column had very clear anatomical and geometric correspondences, in particular within L4. The question however whether the precision of wiring, the relation to column vs septum circuitry, the subtypes of neurons and their higher-order circuitry can also be found in cortices with less explicitly clustered arrangements has to be experimentally determined, in particular for more higher-order cortices.

### Learning: hot-spots of synaptic storage?

The non-uniform distribution of connections with evidence for homosynaptic plasticity points to a concrete site specialized for possible information storage within the cortical column. With the L3-to-L5 connection as the strongest site of synaptic weight similarity, one can speculate whether this corresponds to learned items of matching sensory-related encodings (via L4->L3) to relevant output decisions. In particular, the finding of L1 axons with L3+L5 preference may point to a distally triggered learning of associations in the column. Furthermore, since also the long-range input to L5B seems biased towards stronger signs of learning, one could consider the L5B inputs to be a primary storage site in cortex (Haozhe, Ludovica et al. 2023). Notably, L5 input synapse distributions defied the strong synaptic weight homogenization seen in all other ExN types, further pointing to their possibly unique role in memory storage.

### Outlook

With the first cortical column connectome available, and basic analysis approaches yielding a new level of clarity of the cortical microcircuit, we can now hope to study the degree of modularity across cortices, the integration into thalamo-cortical loops, long-range interactions, and possible modifications of the cortical column connectome across evolutionary time scales. This manuscript reports a template, which future studies can build upon.

## ACKNOWLEDGEMENTS

We are grateful to Axel Borst for pre-publication sharing of the CMSR algorithm, Larry Abbott, Haim Sompolinsky and Axel Borst for fruitful discussions, Ali Karimi, Saksham Pruthi, Ali Hadian and Varun Khare for contributions to data processing and analyses, Iris Wolf, Smaro Soworka, Selina Horn, Lev Dadashev, Thomas Olstinski for experimental support, Christian Guggenberger, Klaus Reuter, Timoteo Colnaghi at the Max Planck Computing and Data Facility (MPCDF) in Garching, and Georgi Tushev, Andreas Kotowicz and Tobias Zeller for computational support, Norman Rzepka, Georg Wiese, Valentin Pinkau and Jonathan Striebel at scalable minds for contributions to automated processing, Matthias Fahland, Nicole Prager and Cindy Steiner at Fraunhofer FEP for providing ATUM tape, Jakob Straehle for contributions to tape optimization, Richard Schalek for initial advice on the ATUM setup, and Heiko Wissler for help with visualizations and management of the student tracer team.

## AUTHOR CONTRIBUTIONS

MH conceived, designed and initiated the study, MSi set up the experimental methodology and conducted all experiments, MSi performed initial analyses and dendritic reconstructions, AM developed methodology and performed automated axon reconstructions, MSch contributed RoboEM technology for axon reconstruction and conducted spine attachment, MH, AM, MSi, MSch performed data analyses with contributions by YY, SL, KS and JB. MH wrote the manuscript with contributions from all authors.

## METHODS

The methods related to tissue preparation, experimental procedures, data processing for 3D alignment and reconstruction are identical to those reported in the PhD thesis of one of the first authors (Sievers 2023). They are reproduced here since they were factually identical to the methods used for this paper. Only minor modifications were applied where necessary. The following methods sections are not put in quotation marks, even though largely text- identical to the methods in (Sievers 2023).

### Animal experiments

All experimental procedures were performed according to the law of animal experimentation issued by the German Federal Government under supervision of the local ethics committees and according to the guidelines of the Max Planck Society, and were approved by the Regierungspräsidium Darmstadt (protocol id: F126/1002). A young adult, male wild-type (C57BL/6J, P28) mouse was subcutaneously injected with general analgesia, 0.1 mg/kg buprenorphine (Buprenovet, Recipharm, France) and 100 mg/kg metamizole (Metamizol WDT, WDT, Germany). Anesthesia was introduced and maintained by inhalation of isoflurane (3 - 4% in carbogen). Next, the animal was transcardially perfused with 15 ml of flushing solution (0.15 M sodium cacodylate, pH = 7.4, Serva, Germany), followed by 30 ml of fixative solution (2.5% paraformaldehyde (Sigma-Aldrich, USA), 1.25% glutaraldehyde (Serva) and 0.5% CaCl_2_ (Sigma-Aldrich) in 0.08 M sodium cacodylate, pH = 7.4) using a syringe pump at a flow rate of 10 ml/min (PHD Ultra, Harvard Apparatus, USA). After perfusion, the animal was decapitated. The brain was exposed by incising a caudo-rostral opening in the skull and immersed overnight in the fixative solution at 4°C.

### Tissue extraction and preparation

After overnight tissue fixation, the sample was extracted from barrel cortex using a 1.5-mm biopsy punch (Standard Biopsy Punch, Integra, USA) mounted on a stereotaxic instrument (David Kopf Instruments, USA). For homogenous staining and embedding, the sample was trimmed to a thickness of 0.4 mm with a vibratome (HM 650V, Thermo Fisher Scientific, USA), resulting in an overall sample size of 1.1 mm^3^ (∼1.5 mm x 2 mm x 0.4 mm in width, height and depth) encompassing all cortical layers plus some subcortical structures. *En-bloc* tissue processing was performed as described in (Karimi, Odenthal et al. 2020). Embedding was based on the Epon hard embedding protocol as described in (Loomba, Straehle et al. 2022).

The positioning of the biopsy punch was determined using a reference atlas (Franklin and Paxinos 2008) and confirmed post-hoc by cytochrome c oxidase staining of the corresponding hemisphere. To this end, 100 µm thick tangential sections were cut with the vibratome and collected into 24-well culture plates. The sections were then rinsed three times in PBS buffer for 20 minutes each. Next, the PBS buffer was replaced by about 1ml of staining solution (0.2 g/ml catalase (Sigma-Aldrich), 0.2 g/ml cytochrome c (Sigma-Aldrich) and 0.5 mg/ml diaminobenzidine (Sigma-Aldrich) in PBS). For better penetration of the staining solution 0.1% Triton-X-100 (Sigma-Aldrich) was added. The slices were incubated in the staining solution at 37°C until the barrels became visible. Finally, stained slices were transferred to PBS buffer, and images were acquired with a conventional stereomicroscope (S6 D stereomicroscope, Leica Microsystems, Germany).

### ATUM-MultiSEM imaging

For ultra-thin sectioning, the cured sample was trimmed to an elongated hexagon shape (EM TRIM2, Leica Microsystems, Germany). A total of 7,199 sections were collected on carbon coated Kapton tape (fabricated in collaboration with Faunhofer FEP, Germany) using a customized ATUMtome setup (RMC Boeckler, USA). Prior to section collection, the tape was plasma treated at a current of 30 mA and a tape advancement speed of about 2.5 mm/s (Q150R ES sputter coater with reel-to-reel tape mechanism, Quorum Technologies, UK). All sections were cut at a nominal thickness of 35 nm (4 mm ultra35° knife, DiATOME, Switzerland) and a cutting speed of 0.3 mm/s within a total experiment time of 28 hours. The cutting direction was aligned parallel to the cortical axis (starting from white matter/subcortical tissue). Two lateral knife shifts were performed, after 120 µm (section 3,434 to 3,435) and 220 µm (section 6,278 to 6,279), respectively. The second knife shift involved a knife cleaning using a polystyrene rod dipped in ethanol. In both cases, the cutting series was resumed after only a single half cut section.

For imaging, the ATUM tape with the collected sections was adhered to 9.5 cm x10 cm silicon wafers (laser-cut from p-doped, one side polished 6’’ wafers, Science Services, Germany) using double-sided adhesive carbon tape (P77819-25, Science Services). On average, one wafer held 225 sections. To get rid of remaining air bubbles and for degassing of the adhesives, the wafers were put in a low vacuum oven (30 mbar, 40°C; Vacutherm, Thermo Fisher Scientific) for a minimum of 12 hours. The wafers were then mounted on dedicated MultiSEM specimen holders equipped with three L-shaped reference points (ZEISS, Germany). An additional layer of carbon tape (P77816, Science Services GmbH) was applied along all edges of the ATUM tape and along all edges of the wafer to enhance conductivity. LM overview images of the wafers were acquired with the ZEISS Axioscope at a pixel size of 3.632 µm (Axio Imager.A2 Vario, Objective: EC Epiplan-Neofluar 5x/0.13 HD DIC M27). Specimen calibration and automated section detection was performed using the Shuttle & Find Module (S&F) and Serial Array Tomography (SAT) module, respectively (ZEN software, ZEISS).

Per section a rectangular region of interest (ROI) with a width of 1.4 mm and a height of 1.5 mm was defined, spanning all cortical tissue. MultiSEM imaging was performed with a 61-beam setup (ZEISS MultiSEM 505; equipped with ZEN MultiSEM software) operated at a pitch size of 12 µm, a pixel size of 4 nm, a landing energy of 1.5 keV and a scan rate per beam of 20 MPx/s. The intra-hexagonal and inter-hexagonal overlap was set to 500 nm and ∼6 µm (10%), respectively. For autoroutines, six support points per ROI were defined. Focus and stigmation corrections were measured at a pixel size of 10 nm and a pixel dwell time of 200 nm. A tilted-plane interpolation was used to derive correction values for the entire ROI. If more than 25 % of the support points failed, ROIs were omitted and revisited in additional experiments. The stage movement and settling time was set to 1 s

All 7,199 ROIs were mapped with the MultiSEM (overall imaged tissue volume: 0.5 mm^3^). Each ROI was padded with 328.9±8.5 hexagons, i.e. 20,069 individual images, whereby the exact tessellation was determined by the orientation of the section. Raw images were saved as 8-bit BMP files, each of which was 8.3 MB in size, resulting in a total raw data amount of 1.1 PB (152 million files). The acquisition time per ROI was 11.9±1.6 min with 33.5 % of the time spent on scanning, 57.7 % on stage movement and 8.6 % on autofocus and autostigmation routines. Additional overhead times were caused by retakes of failed ROIs and wafer exchange. Including retakes, one wafer was routinely imaged within 48 hours, corresponding to an effective imaging throughput of 18 TB/day and a volumetric data rate of 3 mm^3^/year.

### Data handling and image quality monitoring

MultiSEM images were first written to a 30 TB buffer server located at the MPIBR. To monitor image quality, a regular grid of control points with a maximum spacing of 100 µm was defined for each ROI. Per control point, the nearest image was determined and cropped to a size of 512 x 512 pixels. The subimages were then merged into montages that were manually inspected for problematic locations. Monitoring routines were implemented in MATLAB (R2017b) and executed on a local workstation in parallel with image acquisition. For further data processing, the images were transferred to the Max Planck Computing and Data Facility (MPCDF, Garching, Germany) via a direct 10 Gbit/s glass fiber link. Data transfer to the MPCDF was executed in a fully automated manner with custom written bash scripts.

### 3D image alignment

3D image alignment was done based on published routines (Scheffer, Karsh et al. 2013); https://github.com/billkarsh/Alignment\_Projects]. In brief, the alignment consisted of the following main steps: (1) creation of the image database and alignment workspace, (2) extraction of same-layer point pairs, (3) solving of montages, (4) coarse strip-strip alignment, (5) refined block-block alignment, (6) extraction of cross-layer point pairs, (7) solving of the final stack. Creation of the image database in step (1) was replaced by custom written MATLAB (R2017b) scripts. As starting point for the in-plane alignment stitched image positions provided by the MultiSEM ZEN software were used. To build the alignment work space, the minimum image overlap size was set to 50,000 and 500,000 pixels with a xy-confidence value of 1.0 and 0.8, and a minimum correlation value of 0.2 and 0.1 for the same-layer and cross-layer case, respectively. Image overlaps that did not meet these requirements were ignored. Same-layer and cross-layer point pairs in step (2) and (6) were extracted from overlapping regions at original image scale, i.e. 4 nm pixel size. To this end, overlap regions were carved into a mesh of triangles, which were then cross-correlated between images. Cross-correlation was optimized using a deformable mesh algorithm. If the final correlation was above 0.3 and 0.15 for the same-layer and cross-layer case, respectively, the centroids of the triangles were accepted as matched point pairs. The maximum single triangle area change was set to 35% and the maximum summed triangle area change to 10%.

Strip-strip and block-block matching, step (4-5), were performed at 80-fold scale reduction. Being the starting point for all further cross-layer steps, results of the strip- strip matching were manually screened by applying the global affine transformation to the low-resolution montages generated in step (3) using custom written MATLAB (R2017b) scripts. The strip-width was set to 15 tiles by default and was iteratively adjusted if misalignment was detected. For the block-block matching in step (5), a block size of 8 x 8 tiles was used with an xy-confidence of 0.5, a minimum block match correlation coefficient R of 0.2 and a maximum z-index span of 5 sections (relevant e.g. in case of sectioning artifacts, tape damage etc.). In the last step of the alignment routine, step (7), an iterative, global least-square algorithm was applied considering all matched same-layer and cross-layer point pairs (90k least square iterations in total), outputting one affine transformation per individual image. Custom written MATLAB (2017b) scripts were then used to apply the affine transformations, convert the 2D images into 3D webKNOSSOS wrapper (WKW), each containing 1024^3^ voxels (Boergens, Berning et al. 2017).

For image alignment high-performance compute clusters (GABA [https://docs.mpcdf.mpg.de/doc/computing/clusters/systems/Brain_Research.html] and Cobra [https://www.mpcdf.mpg.de/services/supercomputing/cobra]) were used. Jobs for extraction of same-layer and cross-layer point pairs, originally submitted per block, were grouped into task arrays so that only a single job was submitted per section to reduce the load on the SLURM scheduler by two orders of magnitude. As an additional change to the C++ source code, the generation of log files per image pair was removed as this caused too many small files; instead, a single log file per block was written listing all failed pairs and the associated error message for debugging.

Note that 12/7,199 of the imaged ROIs were omitted from the alignment due to large-scale artifacts (>50 % tissue damage). In one case, two consecutive sections were affected, resulting in a gap of 70 nm. Most of the omitted sections (9/12) were related to an ATUM tape damage.

### Nucleus detection and classification

For detection of nuclei, a 3D U-Net (Ronneberger, Fischer et al. 2015, Çiçek, Abdulkadir et al. 2016) was trained on low resolution data (mag 32, i.e. 128 x 128 x 140 nm^3^ voxel size) to predict voxel-wise probabilities for three classes: nucleus, blood vessel and background. Type probabilities were then converted into nuclei and blood vessel segments by applying a threshold of 50% to the respective probabilities and calculating the connected components. Best results were achieved at a size threshold of 26,106 voxels at a precision of 99.7% ad a recall of 99.2%, respectively (evaluation based on manual ground truth annotations of 4,432 somata within a layer 4 centered bounding box).

To classify the detected cell bodies as neurons, glia, or blood vessel cells, a multiclass decision tree ensemble was implemented in MATLAB (R2017b) trained on a set of 18 features: volume and surface area of the nucleus segment (1-2), diameter of the equivalent sized sphere (3), width, height, depth and volume of the smallest bounding box containing the nucleus segment (4-7), volume of the smallest convex polygon containing the nucleus segment (8), ratio of the nucleus volume to the bounding box volume (9), ratio of the nucleus volume to the convex volume (10), length of the three major axes of the nucleus segment (11-13), median of all intensity values contained within the nucleus segment (14), median distance to other nucleus segments (15), median, 0.25 quantile and 0.75 quantile of distance to blood vessel segments (16-18).. On a test set of 125 cells, the decision tree ensemble achieved 100% precision and 100% recall for neurons, 100% precision and 96% precision for glia cells and 94% precision and 100% precision for blood vessel cells.

Note that within the column bounding box, detected cell bodies and assigned types were manually proofread. In particular, border cases were added that were missed by the automation due to the implemented nucleus size threshold.

### Version-1 segmentation

A variant of 3D U-Net (Ronneberger, Fischer et al. 2015, Çiçek, Abdulkadir et al. 2016)was trained to predict voxel-wise nearest neighbor affinities from 3D EM data at a voxel size of 8 x 8 x 35 nm^3^. To train and validate the model, 78 (1 µm)^3^ boxes were randomly distributed within a layer 4 centered subvolume and manually annotated. The boxes were randomly divided into training (80 %) and validation (20 %) set. After 60 epochs of training the configuration with the best performance on the validation set was selected as the final model, which was then applied to the column bounding box. Subsequently, using the predicted voxel affinities, a watershed-based over-segmentation was applied. In a final step, a region adjacency graph (RAG) was constructed, with each node corresponding to a segment in the oversegmentation and each edge corresponding to an interface between segments, and a median-based hierarchical agglomeration was applied as described in (Funke, Tschopp et al. 2019)]. Blood vessel segments were omitted from agglomeration to reduce percolation of mergers. To build the blood vessel mask, predicted blood vessel probabilities were thresholded at 50%. Connected blood vessel components larger than 87,000 voxels were manually proof-read and false-positive components were discarded. The remaining blood vessel components were dilated by 500 nm via a distance transform and mapped onto segments in the over-segmentation. Segments were blacklisted if most of their volume was enclosed by the blood vessel mask.

### Version-2 segmentation

For v2 of the column segmentation, a 3D U-Net for prediction of nearest neighbor affinities with the auxiliary task of predicting long-range affinities was applied as in (Lee, Zung et al. 2017) with minor modifications. Compared to v1, the amount of the manually annotated (1 µm)^3^ training boxes was increased to 410. The additional boxes were randomly distributed within the entire column bounding box. Manually annotated segments were dilated by up to 35 nm. After prediction of voxel affinities, a watershed-based over-segmentation was applied similar to v1, with the following two adjustments: (1) The affinity threshold to define border voxels in the first steps of the watershed procedure was globally set to 50 %. (2) A minimum segment size of 100 voxels was introduced. Smaller segments were removed and filled up by a 2nd watershed run. The subsequent hierarchical agglomeration with blood vessel masking was complemented by a mutual soma exclusion and type restriction. For mutual soma exclusion, based on manually proof-read soma locations within the column bounding box each segment was assigned a unique soma ID. Segments which did not contain any soma node were labeled with soma ID zero. During agglomeration, the soma IDs of the segments were propagated to the agglomerates. Edges were filtered in descending order of affinity and rejected, if they would merge agglomerates with non-zero and differing soma IDs.

The mutual type exclusion was based on voxel-wise predicted neurite type probabilities, which were pooled over the segments in the over-segmentation (see below). Agglomerate type probabilities were then derived as the volume-weighted average of the underlying segment type probabilities. Edges were rejected if both agglomerates were larger than 2 µm^3^ (to be sufficiently confident on the type scores) AND the maximum type probability of the agglomerate was greater than 60% AND the predicted agglomerate types were contradicting. Note: Spine head scores were merged into the dendrite class. The ‘other’ type was treated as unknown and essentially removed by re-normalizing the remaining astrocyte, axon and dendrite probabilities.

### Segment statistics and agglomerate file

For each segment in the over-segmentation, the maximum distance to the segment border, the position (the point with maximum distance to the segment border), the segment volume, the center of mass and the covariance matrix were calculated. The extracted values were sorted by segment IDs and stored in an HDF5 file, called the segment statistics file. The fraction of applied edges in the spanning tree induced by the hierarchical agglomeration was referred to as ‘mapping’ (e.g., 85% of edges were applied in mapping 85). Each mapping also corresponded to a global minimum affinity threshold, with the value depending on the applied segmentation. The hierarchical agglomeration was run for a range of different mappings. Per mapping, the resulting agglomerates were stored as skeleton graphs (consisting of nodes with associated positions in three-dimensional space and undirected edges between these nodes), within a dedicated HDF5 file, called the agglomerate file.

### Neurite type predictions

Analogous to the affinity models, a 3D variant of the U-Net was implemented and trained to voxel-wise predict probabilities for five classes (axon, dendrite, spine head, astrocyte and other, e.g. blood vessel or soma) from aligned and cubed raw EM data at a voxel size of 8 x 8 x 35 nm^3^. A type annotated version of the segmentation ground-truth data, consisting of a total of 69 (1µm)^3^ boxes, served as training data. The voxel-wise predicted type scores were pooled over the segments from the over-segmentation. Per segment, summary statistics (average, maximum and standard deviation per neurite type) were extracted and written to an HDF5 file sorted by segment ID, called the segment types file. The type scores were also pooled over agglomerates and appended as additional dataset to the agglomerate file.

### Soma-based dendrite reconstruction

Soma-based dendrite reconstructions within the column volume were based on v1 segmentation at mapping 65. Soma locations were obtained from the automated nuclei detection and manually proof-read in webKnossos. New trees were added in webKnossos, if nuclei were missed. For proof-reading, EM data were overlaid with a volume map of all agglomerates at mapping 65. If the soma-seeded agglomerate stopped at the nucleus border, an additional node, referred to as support node, was placed within the cytoplasm. In rare cases multiple support nodes were required. Based on the positions of soma and support nodes, segment IDs were determined and mapped onto agglomerate IDs using the agglomerate file. If there were support nodes, the respective agglomerates were combined into a new agglomerate with ID_new_ = max(IDs) + 1 by adding edges from the full region adjacency graph (RAG). Subsequently, non-unique soma-seeded agglomerates, i.e. agglomerates containing multiple soma nodes, were separated as follows: First, two soma seeds were randomly selected within the merged agglomerate. Second, the segment IDs of the two seed nodes were mapped to nucleus IDs (from the automated nuclei detection). If the nuclei components did not cover any other soma location, the nuclei were ‘anchored’ to the agglomerate by setting the affinity of all edges connecting the nucleus segments to the surrounding segments to 255. This step was required to not split off the nucleus in the following steps. Third, the affinity threshold at which the two seed nodes were in separate agglomerates was determined. Finally, edges below the affinity threshold, if part of the current shortest path connecting the two soma seeds, were iteratively deleted, until the seed nodes were no longer connected. The four steps were repeated until all soma seeds were in separate components. New agglomerates with were defined as the connected components containing the soma seeds and appended to the agglomerate file. Corresponding segment IDs were remapped. The agglomerates modified in this way served as starting point for further neurite reconstructions.

Remaining splits in the dendrites of all soma-seeded agglomerates, previously labeled as neurons, were solved using an iterative, semi-automated approach inspired by(Motta, Berning et al. 2019). Endings of the soma-seeded agglomerates were detected as follows: Per node, the agglomerate was restricted to an outer sphere of r = 25 µm, centered on the node of interest (highlighted in red). Only the connected component containing the current node-of-interest was further considered. All nodes within an inner sphere of r = 5 µm (including the node-of-interest) were discarded. Finally, the remaining connected components were counted, corresponding to the number n of inner sphere exits. If n_exit_ = 1, the node was labeled as ending node. All nodes with n_exit_ = 1 and a pair-wise distance of less than 2.5 µm were combined into one ending.

For each ending, the minimum spanning tree (MST) was determined based on the inter-node distances in µm. For focused ending queries using the webKnossos flight mode (Boergens, Berning et al. 2017), the node with the minimum average distance to all other nodes on the MST was defined as start position. The start direction was defined as the unit vector from the start position to the volumetric center of mass of the 90th percentile of all nodes per ending that were more distant from the soma than the current start position. If any of the ending nodes was closer than 1 µm to the border of the column bounding box, the ending was considered as EOD (end-of-dataset) and was not processed further. Endings were also omitted from flight queries if the average type scores for the glia, axon or other class were above 75%. Ending queries were performed by student annotators, who were instructed via a video-based task description. A volume map of all agglomerates larger than 7 µm was used to dynamically stop flight tracings, when more than 10 nodes were placed inside another agglomerate. The agglomerate size was approximated by the diagonal of the smallest cuboid containing the agglomerate. In addition, hotkeys were implemented to indicate true dendritic endings and ambiguous/unclear locations. Both hotkeys immediately terminated the current query. When a query was finished, a new task was started automatically. If more than 400 nodes were placed without any of the above end scenarios occurring, the query timed out.

The ending queries yielded linear skeletons with labels for new agglomerate IDs, true endings, mistracings or timeouts. In the first two cases, the nodes of the flight path were mapped onto the underlying segment and agglomerate IDs, respectively. For true endings, if nodes were placed outside the base agglomerate, the agglomerate graph was augmented with the additional flight nodes and edges. If a new agglomerate ≥7 µm was picked-up by the flight trajectory, the new agglomerate was loaded from the agglomerate file and attached to the base agglomerate via flight nodes and edges (restricted to the shortest path between the two agglomerates). New agglomerate IDs were accepted if they did not belong to another soma seeded agglomerate AND the agglomerate type was neither axon nor glia. In a last step, endings were detected for each accepted agglomerate and new flight queries were generated, using the same routine and parameters as described above. In case of mistracings, timeouts or other problems, the queries were repeated once again.

Custom-written MATLAB scripts (R2017b) were used for preprocessing routines, ending detection, query generation, and query processing. Thresholds were set heuristically. To grow branches iteratively and independently per agglomerate, each soma-seeded agglomerate was stored as skeleton graph in a dedicated MAT file that was updated after each round of ending queries and also contained status information about the detected endings and the processed flight queries. If required, up to six query iterations were performed for each soma-seeded agglomerate.

### Axon branch detection

To detect the axon branch of the soma-seeded agglomerates, the following approach was used: First, per node, the shortest path to the soma d_soma_ was determined using the skeleton representation of the agglomerate graph with edge weights equal to the Euclidean distance in µm. All nodes with d_soma_ ≤ 25 µm were discarded. Next, connected components were computed, ignoring components with less than 30 nodes. For each component, the axon probability was derived from the volume-weighted mean of the corresponding segment scores. If the axon probability was above 24%, the component was labeled as axon branch. All cases with n_axon_ ≠1 were manually corrected in webKnossos. The axon branch detection was implemented using custom written MATLAB (R2017b) scripts. It was performed for all soma-seeded agglomerates based on the v1 segmentation after the semi-automated dendrite reconstruction. The information on the detected axon branches was stored for each skeleton in the respective MAT file by adding binary axon labels for each node/segment, and was used as starting point for targeted axon reconstructions.

### Simplified skeleton graph

To simplify a skeleton, the skeleton was first turned into the shortest-path tree, with the root being a user-specified center node (typically located in the soma of the neuron). The edges of this tree were oriented towards the root and weighted by the Euclidean distance between the corresponding nodes. This tree was then iteratively simplified by the following greedy algorithm until all remaining branches exceeded the user-specified minimal branch length: First, the distance to the first branch node was computed for each terminal (leaf) node. Furthermore, for each branch node, the terminal node with maximal distance was determined. Terminal nodes were then processed in order of ascending distance to first branch node. If the distance from the terminal node to the first branch node exceeded the user-specified minimal branch length or if the terminal node was also the terminal node with maximal distance from the first branch node, then the branch was kept. Otherwise, the branch (the path from the terminal node to the first branch node) was removed. This procedure was repeated until no more branches could be removed.

### Transfer of dendrites to v2 segmentation and sanity checks

Soma-seeded dendrite reconstruction were transferred to v2 segmentation, using mapping 85 with mutual soma and type exclusion as starting point. Prior to the transfer, implausible flight trajectories from the previous ending queries and their corresponding picked-up agglomerates were removed by applying a maximum path and edge length threshold of 50 µm and 10 µm, respectively. Next, all nodes from the reconstructed agglomerate graph, referred to in the following as seed nodes, were mapped onto the new segment and agglomerate IDs. A picked-up agglomerate was only accepted if it contained more than 10 seed nodes, its dendrite probability was larger than 30% and its volume was larger than 1.12 µm^3^ (same threshold as for the spine head attachment; see next section), OR the respective agglomerate contained the soma seed. All accepted agglomerates were then loaded from the agglomerate file and combined into a new agglomerate by adding seed nodes and edges from the old agglomerate graph. The ID of the agglomerate containing the soma seed was propagated to the entire agglomerate.

Segments of the newly combined agglomerate, which were farther than 1 µm from a segment picked-up directly by a seed node from the old agglomerate graph, were labeled as merger and grouped into connected components (using edges from the new agglomerate graph). Mergers smaller than 7 µm were kept (most likely spines; the size was approximated by the diagonal of the smallest cuboid encompassing the merger component). Mergers with more than 20,000 segments were rejected for computational time reasons. The other mergers were more closely inspected and rejected, if the mean dendrite probability was below 30%, OR the covered seed volume was below 5 µm, OR the merger got disconnected at mapping 80, OR more than 10 contained segments overlapped with any other soma-seeded agglomerate. Segments of rejected mergers were deleted and connected components were determined. The connected component containing the soma seed was defined as the new agglomerate.

After transfer of the soma-seeded agglomerates to the new segmentation, the directional consistency of the dendritic branches was checked. To this end, the new agglomerate graph was first converted into a simplified skeleton (minimum branch length = 25 µm). For each node of the simplified skeleton a direction vector v_p_ (unit: µm) was determined according to v_p_ = (x_p_, y_p_, z_p_) - (x_q_, y_q, z_q), with Q defined as the first node along the shortest path from P to the soma at a minimum distance to P of 10 µm. Direction vectors were normalized and the direction change per node P, referred to as θ_p_, was calculated. The direction vector of the most proximal node on the shortest path to the soma was taken as reference. (Note: Direction vectors and angles were only computed for nodes with d_soma_ ≥ 25 µm.) Nodes of the simplified skeleton were defined as critical, if the determined direction change θ_p_ was larger than 85°. If critical nodes could be traced back to a change in the underlying agglomerate ID within a search radius of 10-20 µm, the corresponding agglomerates were deleted from the full agglomerate graph. Only the connected component containing the soma seed was kept.

As additional sanity check, previously detected and connected ‘mergers’, were treated as separate agglomerates, i.e. they were assigned an artificial agglomerate ID. The threshold for critical angles was increased to 110°, if agglomerates were attached within 2.5 µm distance to the main apical path (= shortest path between the apical tip to the soma). Agglomerates, containing nodes of the main apical branch, were generally excluded from deletion. Remaining errors in the dendrites, if detected during visual inspection of the reconstructed agglomerates, were manually corrected by hard-coding a list of agglomerate IDs to be deleted or not to be deleted. To transfer dendrite reconstructions and perform sanity checks custom written MATLAB (R2017b) scripts were used. For spine head attachment and follow-up connectomic analysis reconstructed agglomerate graphs were mapped backed into the agglomerate file format, maintaining the IDs of the original soma-seeded agglomerates.

### Spine head detection and attachment

For spine head detection, a threshold of 50% was applied to the voxel-wise predicted spine head probabilities and connected components were determined. To separate mergers, the spine head components were intersected with agglomerates (v2 segmentation, mapping 85 with mutual soma and type exclusion). Spine heads were attached to the dendritic shaft as described in (Schmidt, Motta et al. 2024). In brief, spine heads were considered attached if they were part of large enough dendrite agglomerate (>500k voxel in mag 2-2-1 = 1.12 µm^3^, ≥80% average dendrite score). For this, predicted spine head positions were mapped onto segment and agglomerate ids at mapping 85 (with mutual soma and type exclusion, without considering previous dendrite reconstructions). Spine heads which were part of large enough agglomerates with dendrite scores <80% were considered as false positives, and omitted from further steps. Spine heads, which were not part of large enough agglomerates were subjected to a two-step procedure: In a first step, spine head positions were checked in iteratively higher mappings (90, 95, 96) by extending the spine head agglomerates up to a radius of 10 µm and checking the underlying mapping 85 agglomerates. Once there was a connection in a higher mapping to exactly one large enough agglomerate with ≥80% dendrite score (evaluated at mapping 85), the shortest paths between the mapping 85 spine head agglomerate to the mapping 85 dendrite agglomerate in the higher mapping was computed and returned for attachment. In a second step, the remaining detached spine heads were traced with RoboEM. Starting directions for RoboEM were identified by searching for directions with low variance across Monte-Carlo dropout samples. Per spine head, the two best guesses were considered and traced until one (or more) stopping criteria were met: (a) a path length of 7 µm was traced by RoboEM, (b) a path length of 200 nm was traced within an agglomerate >500k voxel, and/or (c) the agglomeration bounding box was left. Tracings were accepted and ‘patched in’ when RoboEM found a large enough agglomerate (case b) with dendrite score ≥ 50%. The best guess direction was checked first.

### Layer 1 dendrite reconstruction

Dendrites from layer 1 neurons with cell bodies within the defined column box (n=64) were manually skeleton reconstructed by student annotators with two-fold redundancy. Locations with differing tracings were reviewed by experts. The combined skeleton nodes from both tracings were then mapped onto segment and agglomerate ids. Picked-up agglomerates were rejected if they contained less than 10 skeleton nodes to exclude false-positives due to misplaced nodes. The agglomerate picked up by the soma position was defined as the base agglomerate. Other picked-up agglomerates were attached to the base agglomerate via nodes and edges from the manual skeleton reconstructions (restricted to the shortest path). Similar to the previously described dendrite transfer, segments of the combined agglomerate that were farther than 1 µm from a segment picked-up by the manual skeleton reconstructions were labeled as mergers and grouped into connected components. Merger components smaller than 5 µm and closer than 25 µm to the soma were kept, all others were rejected and removed from the agglomerate.

### RoboEM with local realignment

To resolve split errors at neurite discontinuities caused by misaligned image data, a local realignment component was added to RoboEM. To this end, slice-to-slice shift vectors from 512 × 512 at 8 × 8 nm^2^ pixel size were precomputed on a grid without overlap. Specifically, global maxima of auto- and cross-correlations of mean- subtracted and maximum intensity normalized slices (*z*_*i*_, *z*_*i*_), (*z*_*i*_, *z*_*i*+1_) and (*z*_*i*_, *z*_*i*+2_) were determined corresponding to pixel-level shift vectors 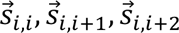. In addition, the mean and standard deviation of correlation-weighted shift vectors reaching a correlation of at least two-thirds of the maximum correlation relative to the noise level were computed. Specifically, shift vectors were considered if the correlation exceeded 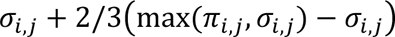, where the peak cross-correlation between slices *i*, *j* is *π*_*i,j*_, and the noise level 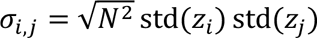 (i.e., standard error of the correlation expected under the assumption of independently and identically distributed pixel intensity values within each slice). To filter irregular shift-vectors, a number of criteria were checked. For the auto-correlation of slice *z*_*i*_ to be considered valid the following conditions were required: 1) auto-correlation must peak at zero shift, i.e., 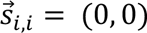, and 2) must be sharp enough with a standard deviation of ≤ 2 pixels, and 3) the summed peak auto-correlation *π*_*i*_ normalized with the expected auto-correlation value *e*_*i*_ = *N*^2^Var(*z*_*i*_) from two successive slices *z*_*i*_ and *z*_*i*+1_ subtracted from the same metric of surrounding three slices *z*_*i*–2_, *z*_*i*–1_ and *z*_*i*+2_, must be ≥ −0.095. For shift vectors 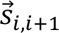 from the cross-correlation of slices (*z*_*i*_, *z*_*i*+1_) to be valid the following conditions were used: 1) auto-correlation must be valid (see above) for both slices 2) the peak cross-correlation normalized with *e*_*i,j*_ = *N*^2^std(*z*_*i*_)std(*z*_*j*_) must be ≥ 0.18, 3) the shift vector standard deviation must be ≤ 10 pixels. For locations, for which 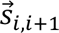 are invalid, shift vectors 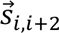 from (*z*_*i*_, *z*_*i*+2_) slices were considered provided that the following criteria were met: 1) auto-correlation conditions (see above) must be true for both slices and 2) the shift vector standard deviation must be ≤ 11 pixels. To compute otherwise invalid shift vectors 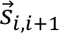, valid shift vectors 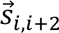 and/or 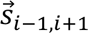 were used. For instance, given valid shift vectors 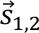 and 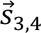, and at least one valid shift vector 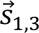 and/or 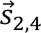, the invalid shift vector 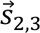 was computed as 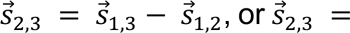 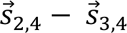, or as the average of those two if both 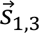 and 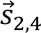 were valid. Similarly, given valid shift vectors 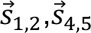, and 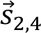, the invalid shift vectors 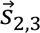 and 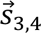 are approximated as 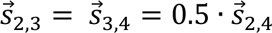. Remaining invalid shift vectors 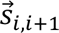 are set to 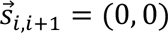.

Within RoboEM, shift vectors were used to generate a locally realigned input and to dealign RoboEM tracings to map back to the coordinate system of the global alignment. To generate the locally realigned and trilinearly interpolated subvolume that is fed to RoboEM, first the current position 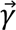 is taken as the reference to compute local shift vectors such that the slice at 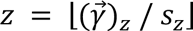 has a shift of 0, where *s*_*z*_ = 35 nm is the voxel size along the slice axis and ⌊*x*⌋ refers to a floor operation. Then, per voxel trilinear interpolation is computed by first linearly interpolating along *x* and *y* axes independently for floored and ceiled *z* values respectively with corresponding shifts along *x* and *y* applied, and then linearly interpolating along the *z* axis. Neurite continuation predictions by RoboEM on the locally realigned subvolume were integrated within the local coordinate system, then aggregated shift vectors were applied to yield the next position within the global coordinate system. For any path length measurements within RoboEM, e.g., used in stopping criteria, the additional path length introduced by the application of shift vectors was ignored.

## VPM axon definition

Similar to (Motta, Berning et al. 2019), thalamic axons from VPM were identified based on the property of establishing large multi-synaptic boutons (Bopp, Holler-Rickauer et al. 2017). Specifically, for excitatory axons with at least 10 spine synapses, spine synapses were clustered to boutons with less than 2.4 µ*m* axonal path length between cluster centers. The subset of axons with at least 50% multi-spine-synaptic boutons and at least an average of 2 spine synapses per bouton were classified as putative VPM axons.

## Axon reconstruction

Axon reconstructions within the column volume were based on v2 segmentation at mapping 85, after transferring soma-based dendrite reconstructions from v1 to v2 segmentation followed by spine attachment. First, one of the neurites exiting the neuronal soma was defined as axon. For pyramidal cells this was possible automatically with proof-reading of the axon definition, investing approx. 50 work hours. For interneurons, axon definition was also attempted automatically first, but required more intense proof-reading.

Then, the axonal neurites were used as seeds for iterated RoboEM reconstruction. For this, the endings of neurites were detected using a method similar to Motta et al 2019. For all detected neurite endings RoboEM was seeded in the direction of the neurite to determine possible axonal continuations. For this, RoboEM would trace until it finds a previously split axon agglomerate to attach to the seed agglomerate. RoboEM was run in validation mode, i.e. a forward and backward RoboEM were performed and only if they agreed the attachment was accepted. Furthermore, additional filter criteria for a successful RoboEM-based axon attachment were employed such as: distance to soma, minimum agglomerate size, and the axon type classification probability; for more details see code repository. After all the seed endings were traced and validated, one round of RoboEM is complete. Multiple rounds of RoboEM were run to iteratively grow axons for each neuron, until there were no new endings detected or for a maximum of 20 rounds.

The implementation criteria were chosen such that the reconstruction was biased towards splits rather than mergers. The degree of completeness of axonal reconstructions was considered in each of the connectomic analyses performed in the study.

After axon tracing by RoboEM, the baseline agglomerate file at mapping 85 with dendrite reconstructions and spine attachment was updated to a newer version with reconstructed axons. During the update, the soma-based agglomerates never change their agglomerate ID. This agglomerate ID was used as “persistent neuron identifier” across further agglomeration versions.

Conflicts in axon attachment are handled while updating the agglomerate file. Segments of the agglomerates that get claimed by multiple agglomerates are removed from all of them. The skeleton nodes themselves are kept. Thus, the morphology remains unchanged. Instead, the conflicting segments are partitioned into separate agglomerates.

Proof-reading for axonal merger corrections was then supported by (1) automatically identifying sections of the axon skeleton lacking the segments for a human expert to decide which neuron gets the contested segments; (2) evaluating morphological features to identify and resolve merges.

For axons that do not originate from a neuron within the dataset, cortical layer one and thalamocortical axons, additional reconstruction was performed. Here a version of RoboEM, that includes local realignment was used. This improved RoboEM version performed better especially in regions with remaining alignment issues particularly in layer one and layer six.

For VPM axons, the largest 500 agglomerates were used as RoboEM seeds to iteratively grow them as described above.

For L1 axons, we followed a different strategy. Approximately 400K agglomerates in L1 was identified as RoboEM seeds. Instead of running iterative rounds of RoboEM on these seeds, only one round of RoboEM was run. After this round, the newly found agglomerates that do not belong to the previous list of seeds were designated as the seeds of the next RoboEM round. This iterative reseeding was repeated for 14 rounds until less than only 0.03% of the found agglomerates were novel. In total, more than 1.5M agglomerates were collected this way. Following RoboEM tracing, an additional consolidation step to identify connected components was necessary before incorporating the agglomerate attachments in an updated agglomerate mapping file, because unlike the soma-seeded axons from different neurons, some of the axon- seeded axons are expected to connect to each other. To reduce merger errors during this process, we excluded certain misclassified agglomerates based on a combination of the following criteria: agglomerate size, number of appearances on the pre- and post-synaptic side of the connectome, excessive number of detected endings, excessive number of RoboEM attachments. This resulted in an approximately 150K connected components that were treated as independent agglomerates for updating the agglomerate mapping as described above.

### ExN clustering (Fig. 2)

The connectome between n=6777 ExNs (based on an earlier tentative ExN definition) was used as the data matrix for hierarchical clustering using Ward’s method (method linkage in MATLAB). Data vectors per neuron were the synaptic inputs to that neuron from all ExNs, reported as number of synapses per connection. The resulting linkage dendrogram was plotted and the first levels of clusters investigated. The first main separation was into L4 of the home column (L4hc) and all other ExNs, followed by a split of home-column L4 into upper and lower L4 (L4hcU and L4hcL, respectively), a separation of L3, L2, L5, and L6, and clusters containing neighboring-column neurons. Notably, L4 of neighboring columns were in separate clusters, each, providing the cortical column (barrel) identification directly from the connectome at the level of layer 4. Furthermore, we noted a home-column aligned L6 cluster, already at this coarse level of clustering (see below). The main layer-related clusters were used to obtain soma position ranges from the percentiles of the soma-depth distributions per cluster (notably without any a-priori depth assumption), which were used in later steps for refining clusters.

We then investigated the L4 home-column cluster (L4hc). In both the upper and the lower L4hc cluster (L4hcU, L4hcL), we noted, in addition to classical spiny stellates, neurons with extended apical-like dendrites (as described for star pyramidals), but notably not a substantial prevalence of pyramidal-like neurons. The fraction of input synapses above L4 (thresholded at 0.5%) yielded a separation into L4hcL-star pyramidals (L4hcLstp) vs spiny stellates (L4hcLsps), and similarly the fraction of input synapses above L3 for the upper L4 cluster (L4hcUsps vs L4hcUstp, thresholded at 0.2%). We then applied a depth homogeneity criterion to remove any remaining outliers: any neuron located more than 4 s.d. from the cluster average were removed. This was <10 out of 400+ for L4hcUsps and L4hcLsps, respectively.

We then investigated the L3 and L2 clusters. The L3 cluster separated into upper and lower L3 (L3A and L3B) clusters, followed by a separation into column aligned clusters and surround. More detailed clustering allowed the separation of L3C clusters. The L2 cluster showed an immediate center-surround structure, which was further separated into L2-only (L2hc and L2sur), and clusters at the same depth as L3A. Together, these clusters were assigned to L2hc, L2sur, L3A, L3B, L3C (hc vs nc was not as pronounced in L3, thus these were kept unseparated at this point).

We then investigated the infragranular cortex. For this, we used the ExN-ExN connectome of roughly the lower half of the cortex (using the whole-column connectome would emphasize intracolumnar translaminar clusters, which were of less interest at this point of the analysis). We used all ExNs at >= 1.75*10^5 voxel depth (corresponding to about middle of L4 and below). The precise position of this boundary did not matter in a wide margin, but we wanted to ensure that that the L4-to-L5A boundary was contained in the lower-column ExN connectome.

Clustering of the lower-column ExN connectome yielded, at the level of soma locations, a separation of L5A, L5B, a home-column centered upper layer 6 cluster (L6Ahc), a less pronounced L6 cluster, and several clusters for the neighboring columns (and the lower part of L4 of all present columns). These are considered the main clusters. Then, the morphologies of all neurons in these clusters were plotted yielding many highly homogeneous clusters, as described in the following.

The main L5A cluster separated into neurons directly below the lower L4 barrel, with dendrites tightly aligned to that lower border (and its notable curvature); and L5A pyramidals without such vertical asymmetry (one of the clusters were large L5B pyramidals which are considered next). The main L5B cluster contained large homogeneous clusters of classical L5Bpyramidal cells, one L5A cluster, and a mixture of L5B and L6, which was assigned later after analysis of the main L6 clusters.

The main L6Ahc cluster separated into clusters with rather homogeneous L6A neurons that extended their dendrite into L4, and more mixed clusters. The other L6 cell types became apparent when analyzing the main L6 cluster, in which a large and homogeneous cluster of L6B cells was assembled, with dendrites reaching to L5A, but not L4. In addition, clusters with local, L6/5B restricted pyramidal-like neurons were apparent.

Based on this tripartition of ExNs in L6, we used the innervation from L4sps, L4stp, L5A, L5Asb and L3 as connectomic data matrix to define these L6 neuron types: first, we determined the mean and variance in this connectomic data matrix of the identified L6A and L6B clusters, respectively. Then, all other clusters were further split according to the cluster linkage sequence, until the cluster members were either at least 75% within the mean+- 1.5 s.d. Euclidean distance from either of the L6A or L6B clusters, or with more than 80% in neither of these spheres, or less than 4 members. In a final step, all clusters with less than 5 members were combined into one, and split by distance to the two L6 clusters (with again 1.5 s.d. acceptance radius). Finally, a depth homogeneity criterion was applied to identify possible outliers from other layers that were sometimes included (+-1 s.d.). The remaining cluster contained partially cut or nc neurons and was not further assigned to connectomic clusters.

Finally, the main clusters comprising mostly neighboring-column L5+6 neurons, and the cluster containing low-input neurons were investigated. In these clusters, several homogeneous clusters with large unusually sparsely branched L5 pyramidal cells were observed. These corresponded to the low-spine-rate input pyramidal cells. Spine rate was used to pull these from the clusters. For the remaining clusters, the above L6A vs L6B analysis was applied, and the resulting clusters were additional L6Bhc neurons, and L6Anc and L6localHC.

Together, this yielded the subdivision of the ExN population based exclusively on connectomic input data (with minor homogenization by depth, but without any pre- conceived cutoffs or assumptions about somatic depth position) as reported in Fig. 2.

### IN type definition (Fig. 3)

The data matrix used for clustering was the combined output and input connectome of all n=1023 INs times all neurons. Clustering was performed using Ward’s method in MATLAB. The first set of clusters already showed strong intracluster homogeneity: n=42 clusters were investigated that comprised 8 clusters of L4-related INs; 1 cluster of L5B-L4 INs; 1 cluster of L3/4 INs; 3 clusters of L3-related INs with however many “mini” INs, that were defined later (see below); 4 clusters of L2-related INs; 7 clusters of L5A related INs; 1 cluster of L5-L3 INs; 4 clusters of L5B-related INs; 2 clusters of NC-related L4+L3 INs; 1 cluster of L1/L2 related INs; 3 clusters corresponding to the L4 INs in the 3 neighboring columns; 4 clusters of L6-related INs; 1 cluster of L5A+B related INs; 2 clusters with mostly bipolar INs. These clusters were then split by soma innvervation vs distal innervation; enhanced IN targeting; higher i/i+e input rate, VPM innervation. Furthermore, innervation of spiny stellates for L4 INs; and spine rate and i/i+e and total synaptic input for “mini” type which occurred mainly in supragranular clusters were used. Finally, clusters with strong parameter similarity were recombined yielding a final of n=55 clusters. Since for these parameter splits, morphology of INs was taken into account, but not used as explicit splitting criterion, the parameter definition sequence represents a connectomic-only type definition with morphology as additional criterion for parameter decisions.

### cFFI + disinhibition analysis (Fig. 4)

For the example analysis of FF circuits (Fig. 4C-F), an individual L3B ExN was chosen and all presynaptic L3B ExNs determined. Only those ExNs with at least 2 synapses per connection are shown in Fig. 4C. Then, all L3sINs were determined that were both presynaptic to the individual L3B ExN and postsynaptic to the pool of connected L3B ExNs. The number of E->I synapses from L3B ExNs onto these individual L3s INs was determined (Fig. 4C). Similarly, L3d INs were analyzed (Fig. 4D). Then, the cFFI synapses from the presynaptic L3B ExNs, L3sINs and L3dINs converging onto the individual L3B ExN were identified and plotted (Fig. 4E), and their postsynaptic path length to the postsynaptic soma determined, yielding the distance histograms in Fig. 4E. This analysis was repeated for n=20 centrally located postsynaptic L3B ExNs and the numbers of converging cFFI synapses determined. This yielded the e/I balance plot in Fig. 4F reporting the relative numbers of I->E synapses from L3sINs, I->E synapses from L3d INs, each normalized to the number of E->E synapses in the cFFI circuits.

For computing the cFFI overlap matrices (Fig. 4G), for each individual IN of a given type, the input from the ExN population (EI, binarized) was multiplied with the output to the same ExN population (IE, binarized), yielding an n_exn_ x n_exn_ matrix reporting which EE pairs were part of a E->I->E configuration. This matrix was element-wise multiplied with the binarized EE connectivity matrix (Hadamard product), yielding those EE connections that were in cFFI configuration for this particular IN. These yielded the columns of the overlap matrix in Fig. 4G. The row means of this matrix yielded the number of cFFI INs per EE connection, and the fraction of rows with at least one non-zero entry the cFFI coverage fraction for this EE connection. These were reported in Fig. 4H for all IN types for a given EE connection.

For determining which INs contribute substantially to cFFI for a given E-E connection (Fig. 4I), the coverage matrices (as in Fig. 4G) were analyzed for all E-E connection types and all IN types. For those IN types where the cFFI coverage (as in 4H) was above 90% or the highest possible for a given E-E connection, their contribution to cFFI was noted in the matrix in Fig. 4I. For judgement of substantial I-I connections (purple data in Fig. 4I), the number of converging synapses from a given IN type to the individual neurons of a postsynaptic IN type were investigated, and only when this was systematically above 10 synapses, the connection was considered substantial. Examples are shown in Fig. 4J,K.

### L1 input analysis (Fig. 5)

For a first estimate of total L1 input synapses to types of ExNs (Fig. 5A,B), a fixed depth threshold of 76000 voxels along y was used (above the L1-L2 border). For further analyses, the following approach was used to obtain a geometrically less biased set of L1 axons: from the initial agglomeration (v2), agglomerates with at least 10 output synapses and at least 90% of output synapses within L1 (above 80000 voxels in y) and at less than 100 µm distance from the approximate home column center in the tangential plane were identified. These agglomerates were used as starting seeds for RoboEM-based axon reconstruction using local realignment (see above), yielding 130,511 column-related L1 axon agglomerates. L1 axons with at least 50 output synapses were used to search for preferential L2 vs other target-innervation (Fig. 5C). Based on the density of L1 axon preferences (Fig. 5C), axons with at least 2/3 of their output onto L2HC ExNs were identified as L2-preferring L1 axons, the remainder as non-preferring for the following analyses.

### Analysis of coordinated Martinotti innervation (Fig. 5I,J)

For the analysis of possibly coordinated innervation via the axon of Martinotti INs projecting both to infragranular layers and L1, the connectome from Martinotti INs with at least 10 synapses in L1 to ExNs of type L5B was restricted to synapses made in L1 (using 76,000 vx in y as cutoff as above); and for the same Martinotti INs the connectome was computed for synapses made below the middle of L4 (Fig. 5I). Then, the two restricted connectomes (L1 restricted and below L4 restricted, respectively) were element-wise multiplied (Hadamard product) and binarized. Then, the row sums of this product connectome reported the number of jointly innervated ExNs per presynaptic IN. These row sums were sorted and plotted. For comparison, the L4 restricted connectome was shuffled along rows, and the resulting joint random innervation plotted as control. For L5B as target ExN type, the actual connectomes showed slightly higher coordinated innervation than random comparisons, but not for L5A. Joint innervation examples of L5B ExNs innervated in L1 and below L4 by two Martinotti INs, each, are shown in Fig. 5J.

### Flow of information analysis (Figs. 6+7)

The circuit diagram in Fig. 6E is based on the type-type connectomes shown in Figs. 6A-D. These connectomes report the average number of convergent input synapses of the individual neurons of the postsynaptic type. With this, a biophysically realistic readout of connectivity can be evaluated that combines the prevalences of pre- and postsynaptic types and the strength of the connection.

The flow of information propagation (Fig. 7E) was computed as follows. The L1 input to all ExN types was row normalized and multiplied to the EE type connectome (shown in Fig. 7A). The resulting vector was row-normalized and considered iteration 2 of information propagation. This was repeated 4 more times to yield the matrix reported in Fig. 7E, left (with normalization following each multiplication). Similarly, the VPM input vector was used to compute the VPM propagation matrix iteratively, shown in Fig. 7E, right.

### VPM subtypes

First, the automatically detected VPM agglomerates (see above) were selected for those with strong innervation of L4Lsps, and the least strongly L4Lsps innervating, yielding a separation as shown in Fig. 6I. Since the length of automated agglomerates was shorter, however, this separation could not prove the two VPM populations to correspond to separate input axons. To obtain ground truth on this finding, first, large axons were selected from the VPM-associated agglomerates (see above). These were inspected manually and corrected for possible reconstruction errors. Then axons were inspected for possible dichotomy into L4-focused and L3-focused axons. In fact, such a dichotomy could be found (Fig. 6J,K), suggesting VPM to project in at least two separate channels to L4 and supragranular layers (see also (Zhang, Wang et al. 2021)).

### Top-down input analysis (Fig. 8)

For the analysis of possibly sorted vs. random intermixing inputs to ExNs in L1 (Fig. 8H-L), the following approach was used. For a given postsynaptic ExN type, the connectome *C_L1ExN_* between all L1 axons with at least 50 output synapses (n=2300) to all ExNs of that type was computed (Fig. 8I). Then, the pairwise input correlations were computed as *C_L1ExN_^T^ * C_L1ExN_,* with all entries in the upper triangle of the resulting product matrix reporting all pairwise input correlations. The histogram of these is shown in Fig. 8J. For comparison to randomly shuffled matrices, the inputs of the ExNs were shuffled by randomly resorting each column of *C_L1ExN_*. With this cases of very high input numbers would not dominate the effects (compared to random shuffling of axonal outputs). Random shuffling was repeated 10 times (black curves in Fig. 8J ff). Then, the same analysis was repeated for the two populations of L2-preferring and non-preferring L1 axons identified before (see Fig. 5C ff), and various ExN types with dendrites in L1 (Figs. 8K,L).

### Connectome sorting analysis (Fig. 9)

For the analysis of possible feed-forward structure in homotypic E-E connectomes (Fig. 9), we applied the smooth-index sorting algorithm (Borst 2024) using the “feedforward” optimization. This resulted in a sorting of neurons. As control, we generated row-shuffled (output-degree preserving) versions of the connectomes. We also compared to Erdös-Renyi random connectomes with matched pairwise connectivity. Feedforward-fraction is reported as 1-recurrent fraction (see Material and methods in (Borst 2024)).

For statistical testing, we computed the feedback and feedforward connection probability for each neuron in the sorting, and evaluated whether the feedforward connection probability was larger than the feedback connection probability for neurons in the middle of the sorting (25^th^-75^th^ percentile) using paired Wilcoxon signed-rank test.

For analysis of effects onto translaminar connectivity, we used the sorting of the L4Usps connectome to also sort the L4Usps->L3A connectome and reported the summed number of synapses per presynaptic neuron. This curve was low-pass filtered (moving median and moving mean function in matlab using a filter size of 50 and 100), yielding an almost 2-fold increase in L4Usps to L3A connectivity.

### Learning analyses (Figs. 10+11)

The following model for learning in homosynaptic connections (same axon-same dendrite pairs of synapses) was used: Synapse-specific noise was assumed to be multiplicative (based on reports of spine volume-dependent volume fluctuations; see for example Box 3 in (Kasai, Ziv et al. 2021) such that for the synaptic weight *w_*i*_* of synapse *i*:

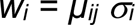

For synapses *i* and *j* from a synaptic pair *ij*, σ_i_ and σ_j_ were assumed to be i.i.d. samples from Lognormal(mu=sigma^2, sigma^2=sigma^2).

Given *w_i_ = µ_ij_* σ*_i_* and *w_j_ = µ_ij_* σ*_j_*, we then derive the maximum likelihood estimate of *µ* and σ (reported as MATLAB formula):

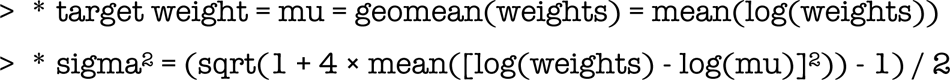

With this, the lower sigma, the closer a connection is to the purely homosynaptic state *w_i_ = w_j_ =µ_ij_*.

The learning analysis can also be described as follows: For each measured and control connection, we can calculate the maximum likelihood estimate of the homosynaptic noise parameter (or the coefficient of variation, or another measure of synapse size similarity). For each value of sigma, we can then ask: What fraction of measured connections have a sigma below that threshold? And what is the fraction of random pairs with a sigma below that threshold? By taking the difference between these two fractions, we can calculate the over-representation of connections below a certain noise threshold. The learned fraction can then be estimated by the maximum difference across all possible thresholds.

So in brief, the learned fraction is calculated as the maximum difference between the cumulative distributions of the noise parameter.

This is identical to the statistic used by Kolmogorov-Smirnov’s one-sided test of equality of continuous distributions. The “learned fraction” is reported only for connections where the distributions of the noise amplitudes of the measured and control connections are significantly different (p < 0.05 after Bonferroni correction).

